# Testing the accuracy of species distribution models based on community science data

**DOI:** 10.1101/2023.01.13.523331

**Authors:** Mélusine Velde, Jacob C. Cooper, Holly Garrod

**Affiliations:** The College at University of Chicago, University of Chicago, Chicago, IL 60637, USA; Negaunee Integrative Research Center, Field Museum, 1400 S Jean Baptiste Point du Sable Lake Shore Drive, Chicago, IL 60605, USA; Committee on Evolutionary Biology, University of Chicago, Chicago, IL 60637, USA; University of Kansas Biodiversity Institute, 1345 Jayhawk Boulevard, Lawrence, KS 66045, USA; Costa Rica Bird Observatories, San José, Costa Rica; BirdsCaribbean, 841 Worchester Street, Natick, MA 01760, USA; Villanova University, 800 Lancaster Avenue, Villanova, PA 19085, USA

## Abstract

While traditional methods of tracking species, collecting specimens, and performing surveys are known to be accurate, additional opportunities to broaden the data pool are evolving. Community science data^5^ has emerged as a new way of gathering large amounts of data, but little research has been done on its reliability for making models for novel locations. The goal of this project was to test the reliability of eBird data as the primary dataset for ecological niche modeling by determining the accuracy of models derived from the citizen-science based eBird dataset. I made species distribution models of 676 bird species in Costa Rica based on eBird observations to predict which species would be found in two localities in Costa Rica that were surveyed. I compared the predictions with these field surveys to determine the prediction success and Sorensen index of the models. Overall, I found that while spatio-temporal factors can affect the accuracy of ecological models, eBird data have great potential as data for species distribution modeling. The models more accurately predicted the community composition in the rural locality as opposed to the more urban locality, and the accuracy of the models increased when compared with data that covered two month as opposed to one month time periods. I tested to see how the number of observations per species influenced the predictive ability of the models and determined that an intermediate number of observations led to better models. These are important metrics to understand because modeling can be an informative and cost effective way to monitor inaccessible areas and can be used in conservation efforts.

## 1. Introduction

Ecological niche models (ENMs) use algorithms to predict suitable conditions for species across geographical space or time based on environmental and ecological factors. Occurrence points are correlated with environmental variables to determine the suitability and consequently predict the distribution of the species in question (Franklin, 2010). ENMs and their derived species distribution models (SDMs) are useful across the field of ecology because they can be applied to most types of living organisms across different types of ecosystems (Elith & Leathwick, 2009). While ENMs predict suitable habitat for a given species, SDMs predict distributions across geographical space to identify their potential range (Peterson & Soberón, 2012). SDMs are an important tool for identifying the ranges of species of interest, such as rare (i.e., Threatened or Endangered) species (Chiffard et al., 2020; Guisan et al., 2006; McCune, 2016) or invasive species (Arenas et al., 2017; Venette et al., 2010). These considerations have important uses in conservation efforts, as they can be used in the selection of conservation sites based on species diversity, species of concern (Loiselle et al., 2003; Moilanen, 2005) or ‘flagship species’ (Cianfrani et al., 2011) present in those areas. They can be key tools used to determine how areas should be prioritized for conservation efforts at the community level (Zhang et al., 2012) or to predict the change in ecosystems as a result of ecological restoration in the face of climate change (Jarvie & Svenning, 2018; Riordan et al., 2018). Species distribution models derived from ecological niche models allow conservationists and scientists to make more informed decisions remotely and with fewer costs; their versatility is making them an increasingly common tool in ecological research.

One of the difficult aspects of deciding which areas should be prioritized for conservation action or exploratory field work is gathering enough data to inform these decisions. When the areas in question are not easily accessible for data collection, it can be difficult to assess the need for conservation efforts or the locations most worthwhile for specific conservation actions (Reddy & Davalos, 2003). In addition, traditional methods of data collection such as tracking species across their ranges tend to be expensive and the training and expertise required to conduct the research is extensive. While these methods are still of immense value for researchers due to their known accuracy (Coxen et al., 2017), community science has emerged as a new form of data collection in the last few decades and has provided researchers access to large amounts of data at a lower cost (Follett & Strezov, 2015). One of the largest such community science projects (especially for birds) is the eBird project (Sullivan et al., 2009).

eBird is a continuously growing community science database that allows birders and ornithologists to report their observations of birds in the form of checklists denoting both presences (i.e. species detected) and inferred absences of species at a given site. eBird checklists are submitted from all over the globe and offer extensive data on the presence of species worldwide that is continually added to and updated. The eBird dataset is free for researchers to download, making it a valuable tool for ornithological studies worldwide (Johnston et al., 2019).

However, there are limitations associated with using community science data. Easily detectable or identifiable species are reported much more frequently than rarer or more elusive species (Sullivan et al., 2009). There is variability in the skill of participants, and misidentifications occur (Bonney et al., 2009). Additionally, temporal biases exist such as the uneven distribution of observations across months within each year (Johnston et al., 2019). From a niche modeling perspective, there are inherent spatial biases that can affect the models. Some checklists are associated with hotspots where observations are spatially concatenated, biasing the data toward specific locations. The locality of the checklist is also a single point as opposed to the entire area that the observer covered. Additionally, most amateur birders tend to stay in easily accessible areas such as near urbans areas, roads, and trails (Johnston et al., 2019). Denser and less accessible areas are neglected and this bias can have significant impacts on where species are reported.

To combat these limitations, eBird provides some quality control for the data. The checklists are required to include certain data relating to effort and completeness of the observations; when a checklist is labeled as “complete”, researchers can assume that species missing from the checklist are absent from that locality. Furthermore, unusual observations are flagged and additional information is requested from the observer; these observations are then assessed for confirmation (Johnston et al., 2019).

While the ornithological research community has begun to use eBird as a primary dataset, little research has been done to determine how reliable those data are for modeling ecological niches. The goal of this project is to assess community science data as a useful tool in conjunction with traditional data collection methods to accurately model community composition. For this project, I made species distribution models using eBird data and compared their predictions to field surveys for two sites in Costa Rica not visited by eBirders to assess the performance of the eBird models.

## 2. Methods

### 2.1 Fieldwork

For field data, I used expert audiovisual surveys and mist-netting data to determine the “true” community composition. I used observation data collected at two sites at different times of the year: Madre Selva in the winter and San José in the summer. I went to Costa Rica in December 2019 with Jacob C. Cooper^6^ and Holly M. Garrod^7^ and we spent 21 hours performing diurnal and nocturnal audio-visual surveys of birds near the Madre Selva research station. We documented as many species as possible with photographs and audio recordings to create a permanent archive of our survey efforts, uploading all material to the Macaulay Library (Parker, 1991). The Madre Selva research station is a privately owned site administered by the Costa Rica Bird Observatories (CRBO) located adjacent to the Pan-American highway at the border between the San José and Cartago provinces (9.661011, −83.867218). We deliberately chose Madre Selva as it had data gaps for the time of year we were there (December). The surveyed area included Isthmian Pacific moist forests at ~2350 m in elevation, and denser Talamancan montane forest at ~2500 m in elevation. In addition to the surveys, we included data from banding operations performed by and in conjunction with CRBO personnel for the months of December and January. At the second locality, surveys were performed in the Montes de Oca canton (9.9554869, −83.9837813) near the city of San José in June and July by Holly M. Garrod. The area is at ~1400m and is primarily Costa Rican seasonal moist forest. All work in Costa Rica was performed with Costa Rica Bird Observatories under their existing permits for ornithological work (including bird banding) in Costa Rica. Jacob C. Cooper and I were covered by an existing Field Museum IACUC under Dr. John M. Bates.

### 2.2 eBird data

Jacob C. Cooper downloaded all of the observations of birds in Costa Rica from the eBird database (version ebd_relAug-2019). The resulting ‘.csv’ file included many pieces of metadata including length of checklist, distance covered, number of observers, etc. I filtered the checklists by effort (< 10 km) and by duration (<500 minutes) to avoid spatial and temporal biases. I then reduced the dataset to unique combinations of species scientific name, latitude, longitude, and date observations. Since I was only modeling the species for the months of December/January (henceforth referred to as ‘winter’) and June/July (referred to as ‘summer’), I created two datasets for each time period. Due to the presence of long-distance and altitudinal migrants, the bird communities of each dataset differ. As elevational migration is documented within resident communities, it is important to only use the observations from the relevant times of year for the most accurate models (Boyle et al., 2016).

I created training areas manually in QGIS for each study locality. The training areas were defined by biogeographical barriers such as mountains, rivers, and valleys (Figure 1). Because of these natural barriers, some species of birds are not physically able to disperse to each area, even if the disjunct habitat is suitable for them (Soberón & Peterson, 2005). To avoid modeling birds that cannot be found at our study site for geographical reasons, I made species lists which only included species that had observations within each training area. In addition, restricting the models with training areas that take into account the species’ known dispersal area increases the predictive ability of stacked (i.e., joint) SDMs (Cooper & Soberón, 2018). For my project, I used the same training areas for all species under the assumption that they all have similar dispersal abilities within the area of study. The models themselves were created over the whole country and the training areas were only used to identify species.

**Figure 1:**
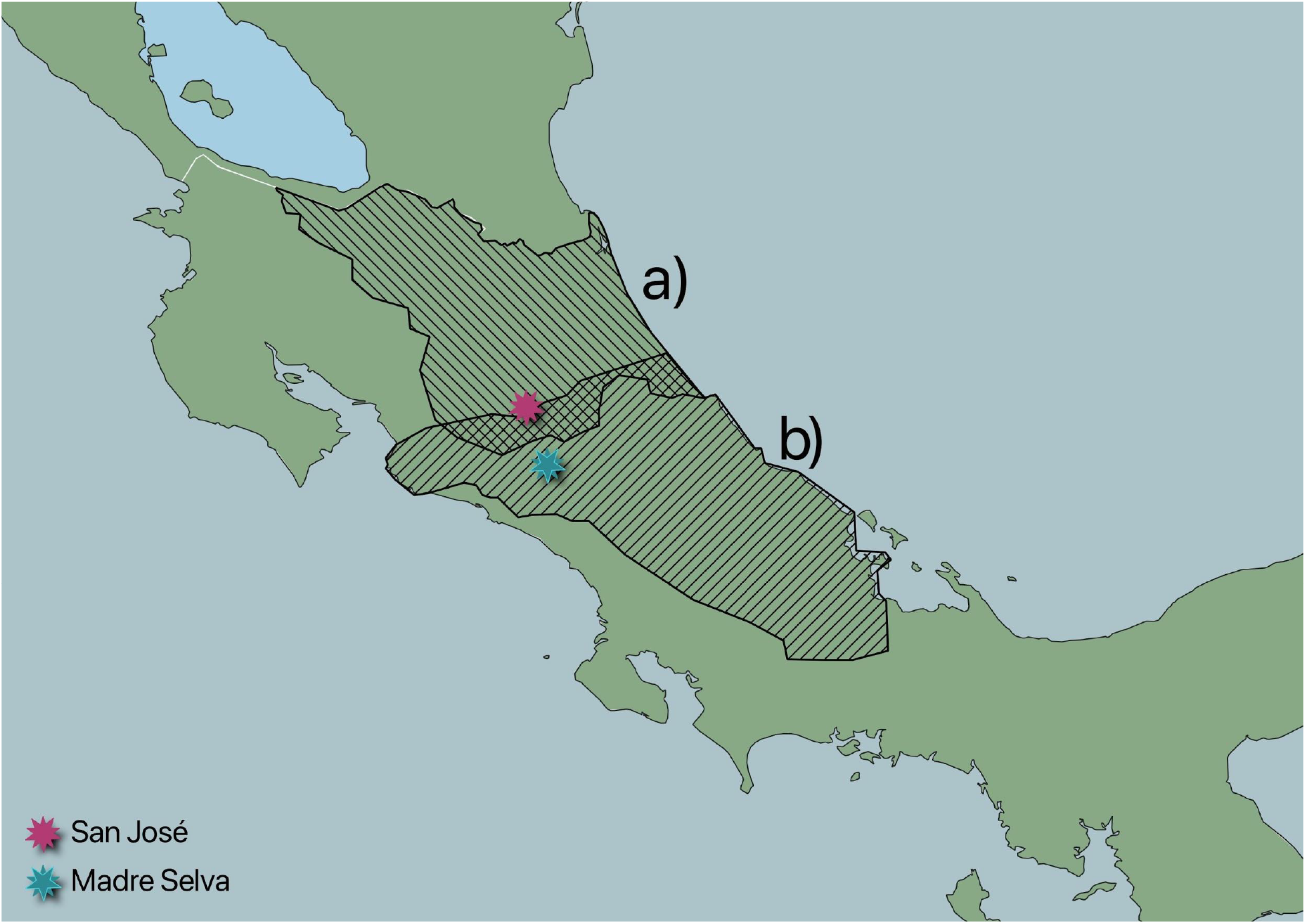
Training areas used to identify the study species for a) San José for the summer b) Madre Selva for the winter

**Figure 2:**
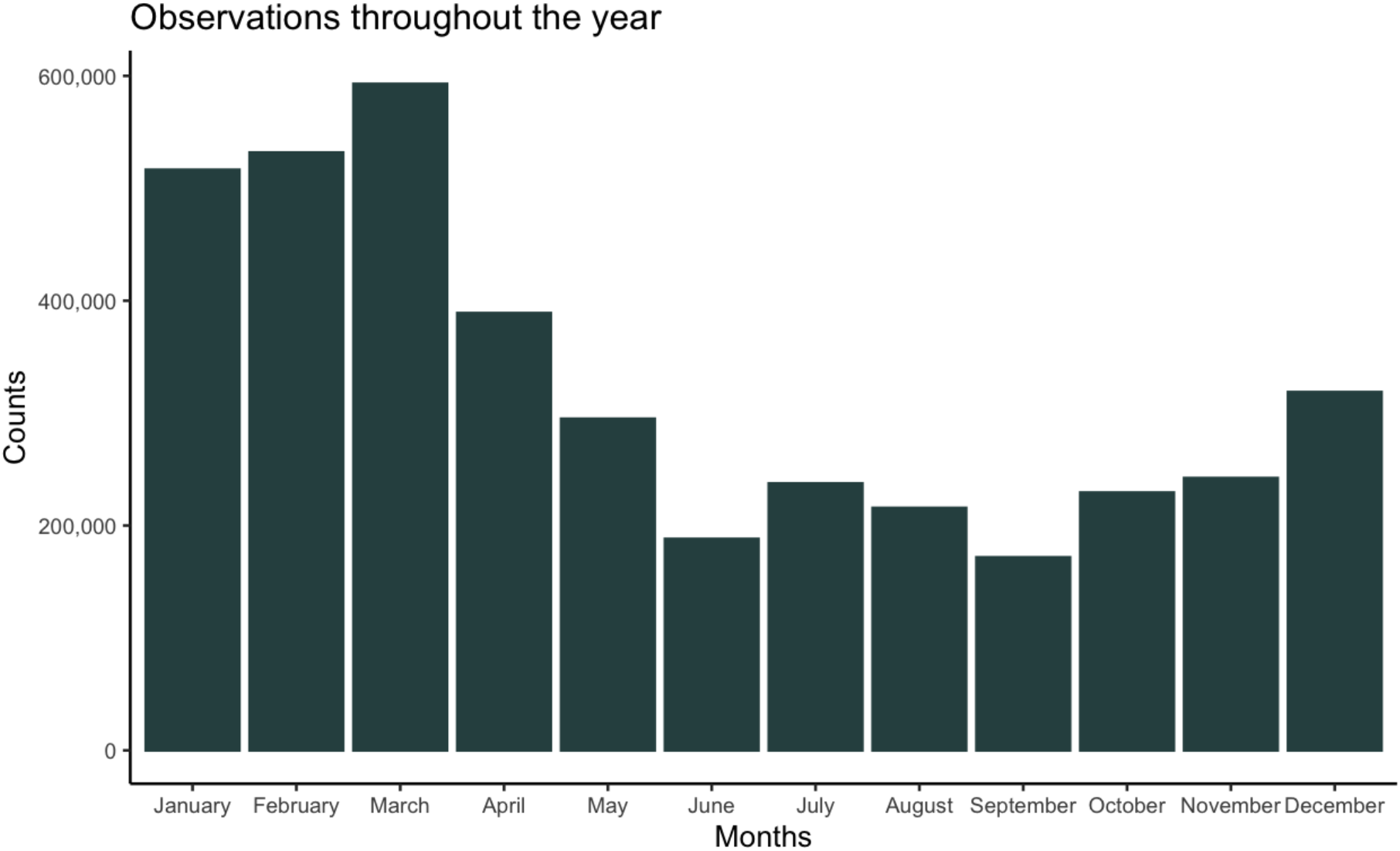
Number of eBird observations by month in Costa Rica and Panama throughout the year (data from 1966-2019)

I spatially thinned the data to unique combinations to remove observations of the same species that were within 1 km of each other. This step was taken to avoid overfitting the models to locations where there are large concentrations of observations. In addition to rarifying, I created an artificial absence of data points within 10 km of the Madre Selva research station and the San José site. Since the goal was to predict which species were found there, I wanted to ensure that there was no spatial bias from imprecision in checklists and to remove the existing data near these sites. Lastly, I removed species with fewer than 10 observations as models were not able to run with fewer data points than the number of datalayers +1.

### 2.3 Species Distribution Models

I used 7 Envirem datalayers (Title & Bemmels, 2018) based on their usage in modeling other tropical montane species, as follows: Thornthwaite aridity index, continentality, embergerQ (a measure of precipitation), maximum coldest temperature, minimum warmest temperature, PET driest quarter, PET wettest quarter (Cooper et al., in press; Title & Burns, 2015). These data layers are relatively uncorrelated and reduce risks of overfitting models in the region (Peterson et al., 2011). As additional datalayers, I used the Normalized Difference Vegetation Index (NDVI) layers which are a measure of greenness in land cover. I obtained these data through the NASA website, where the earthdata page provides open access data that can be easily downloaded (urs.earthdata.nasa.gov). For both time periods I downloaded the MODIS/Terra Vegetation Indices 16-Day L3 Global 1km SIN Grid V006 tiles from 2000 to 2019. Because the month of December is at the transition between the dry and wet season, I used the minimum and maximum NDVI for each year (each averaged over the last 20 years) as two separate datalayers. Using the average minimum and maximum takes into account all seasonal variation and captures the extremes that the birds experience over the course of the year. The datalayers were then stacked and projected onto the study area so that the locality of each observation can be associated with the environmental conditions of the 9 datalayers. The same was done for the month of June.

The ecological niche models were created using minimum-volume ellipsoids. This multi-dimensional ellipsoid is the smallest ellipsoid that encompasses all data points, and it uses Mahalanobis distances to estimate the distance between groups of points (Mahalanobis, 1936; Van Aelst & Rousseeuw, 2009). For a given species, the center of the ellipsoid represents its most suitable habitat based on environmental conditions, and the farther a data point is from the center of the ellipsoid, the less likely it is that the species in question will occur there. I used minimum-volume ellipsoids as they utilize presence-only data, whereas many other species distribution modeling methods require presence/absence or pseudoabsence data (Brotons et al., 2004; Elith et al., 2020). Because my study sites are under-surveyed areas, I have limited absence data and rely entirely on presences. Another advantage is that using this method makes it easy to find outliers as they are at the edges of the ellipsoid (Abou-Moustafa & Ferrie, 2007; Sun & Freund, 2004). To remove the outliers and to create binary prediction models (i.e., SDMs), I used thresholds of 100%, 90%, and 75% data inclusion.

### 2.4 Statistical analysis

Once I had created the SDMs, I made a presence-absence matrix (PAM) for each locality. For each threshold, I identified which species were predicted to be present and absent at the test locality and compared the list to the species actually observed during surveys. For the Madre Selva dataset, I supplemented the data we collected with observations by other researchers in the area for December and January to ensure that the shortcomings in our data collection were accounted for. For the San José dataset, I used Holly M. Garrod’s observations for June and July to compare with the models. This was also a way to test the model’s accuracy. If the models based on a single month perform better when compared to the December + January (‘winter’) dataset rather than just our December data, we can conclude that the models gave a more accurate prediction of the species presence than our own limited-time field surveys. The same is true for the San José data; I compared the June and the June + July (‘summer’) datasets to see which performed better.

From the presence-absence matrices, I calculated the prediction success (*P*; equation 1) and Sørensen index (*S*; equation 2). Prediction success evaluates how many species were predicted correctly out of all the predictions, whereas the Sørensen index weighs presences more heavily than absences. This is important as I do not have true absence data and the Sørensen index takes this bias into account and because presences are more important from a conservation standpoint. These metrics are calculated using the proportion of true positives (tp), true negatives (tn), false positives (fp), and false negatives (fn) as follows:

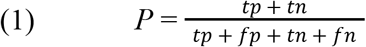

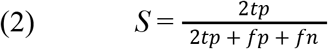

Where:

- <u>tp</u> = the number of species both predicted and observed to be present.

- <u>tn</u> = the number of species predicted to be absent and were not observed.

- <u>fp</u> = the number of species predicted to be present and were not observed.

- <u>fn</u>=the number of species predicted to be absent and that were observed.

### 2.5 Binning by number of observations

One of the major sources of variation across the models was the number of datapoints. While some species were more common or occur in easily accessible areas, others were not as easily detected and consequently less frequently recorded. After reducing and rarefying the data, the range of observations per species was between 10 (the threshold for inclusion) and 1378 (*Coragyps atratus*). Some studies have shown that having too few data points can reduce the accuracy of species distribution models (Wisz et al., 2008) and that these models should be taken as identifying suitable environments rather than distributions (Pearson et al., 2007). On the other hand, reducing the data is important for species with abundant data because too many raw data points can bias the models as well (Ruete et al., 2020). For example, a very common species is more likely to have vagrants that end up in areas that do not match their usual habitat, which could skew the models toward those areas.

I performed an analysis of the reliability of the models based on the number of datapoints. I ordered the species by number of observations and binned them into groups of roughly equal numbers of species. I created a presence absence matrix for each bin to determine the accuracy of the models. I only performed this analysis for the 90% threshold because it had produced the most accurate models according to the Sørensen index. In addition, to compare the summer and winter datasets and their pattern of data collection, I performed a Kolmogorov-Smirnov Test.

### 2.6 Data Availability

All data used in this study are publicly available for research via eBird, including the field data from the two study sites. All aforementioned codes are available in Appendix 1.

## 3. Results

After spatially thinning the data, there were more observations in the winter months than in the summer months, with 835,111 observations in December and January and 425,558 observations in June and July. There was an average of 417,555 observations per month for the winter dataset and 212,779 observations per month for the summer dataset. I modeled 651 species from the winter dataset and 506 species from the summer dataset (Appendix 2). The number of observations per species ranged from 10 to 1378.

### 3.1 eBird datasets

Both datasets have a right-skewed distribution, with many species having few observations (Figure 3). The average number of observations per species was 157 in the winter dataset and 127 in the summer dataset; the median was 92 for the winter dataset and 72 for the summer dataset. According to the Kolmogorov-Smirnov Test (D= 0.096579, *p* = 0.009), we can reject the hypothesis that the two sample distributions are the same.

**Figure 3:**
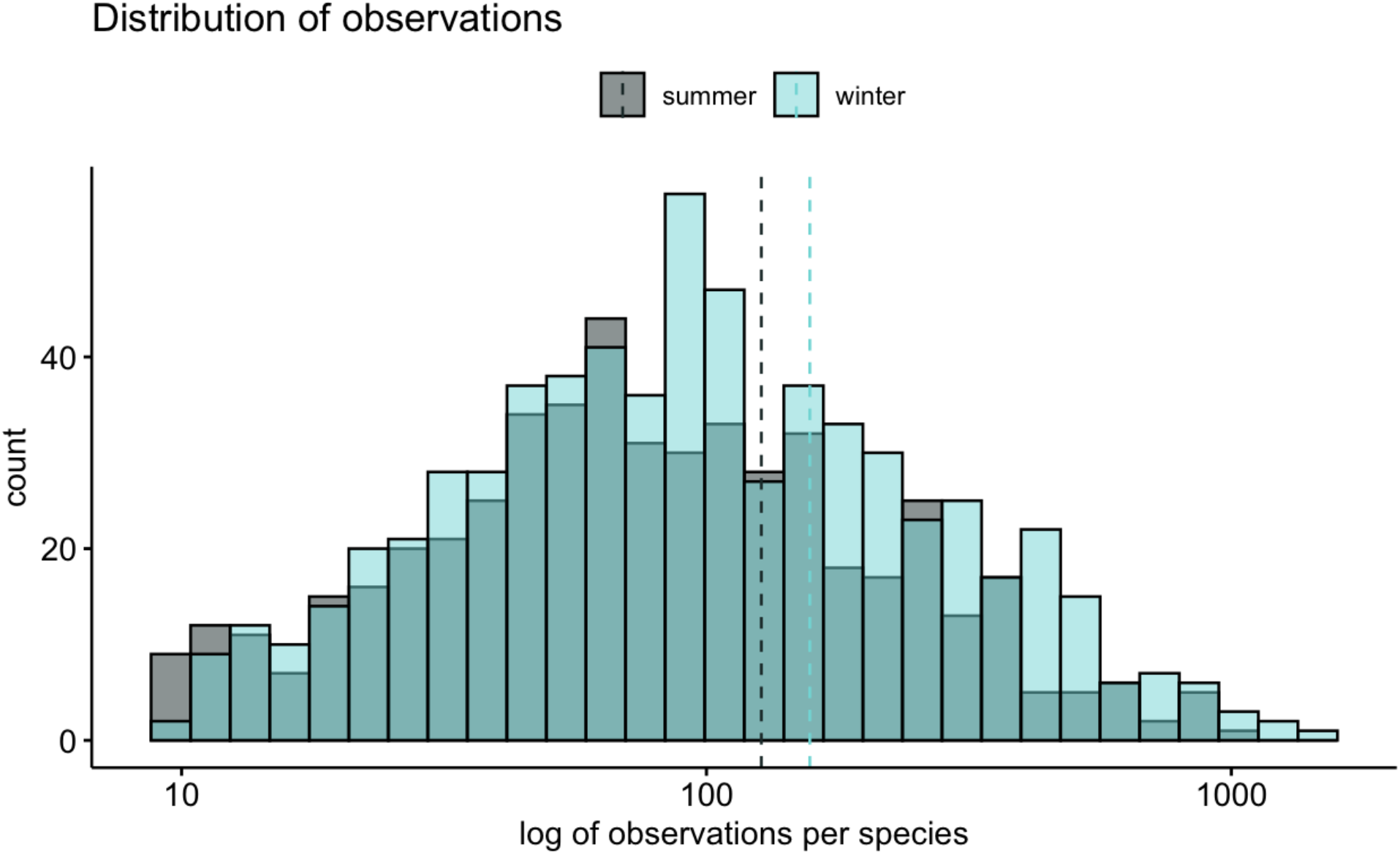
Distribution of summer dataset and winter dataset; dashed lines represent the median for the dataset

### 3.2 SDMs

Four types of SDMs were obtained for each species and an example of each is illustrated for *Myadestes melanops*(Figures 4 & 5). Figure 4 shows the presence absence models are various thresholds (75%, 90%, and 100%) and Figure 5 shows a gradient of suitable to non-suitable habitat based on Mahalanobis distances, or distance from the center of the minimum-volume ellipsoid. The models with the 90% and 100% thresholds (Figures 4e & 4f) show more suitable habitat as more observations were reflective of the distribution while the lower thresholds show a more localized range (Figures 4a & 4b).

**Figure 4:**
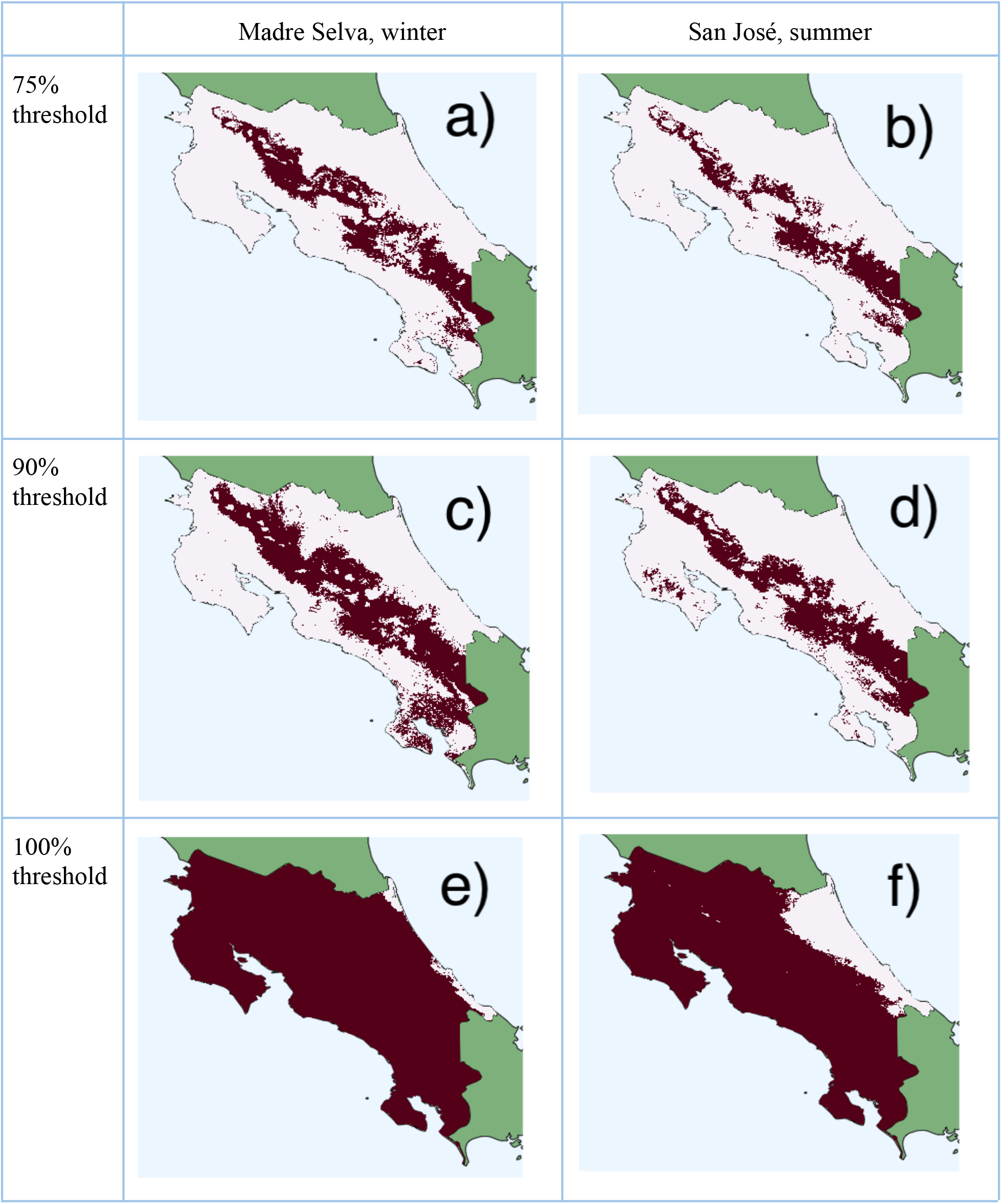
Examples of presence/absence models obtained for the species *Myadestes melanops*; light pink indicates unsuitable habitat, dark pink indicates suitable habitat.

**Figure 5:**
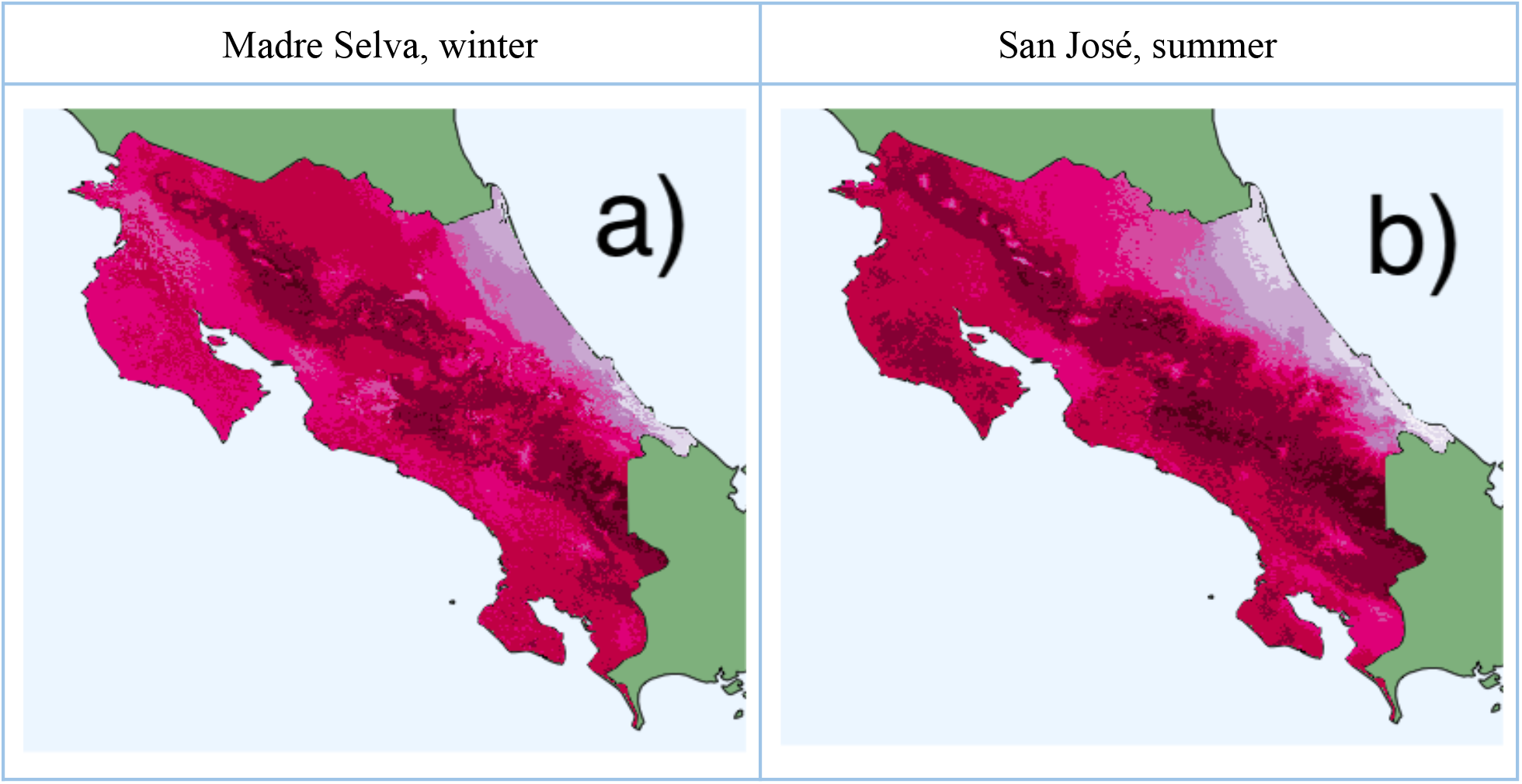
Examples of species distribution models obtained for the species *Myadestes melanops*; the model shows a gradient of most suitable (dark pink) to least suitable habitat.

### 3.3 Comparing one-month vs two-month data

With the exception of the 75% threshold for December (which is higher than the 75% threshold for winter), the two-month datasets were more accurate than the one-month datasets (Figure 6). Hereafter, I present the results for the two-month datasets because they performed the best; the results for the one-month datasets can be found in Appendix 3.

**Figure 6:**
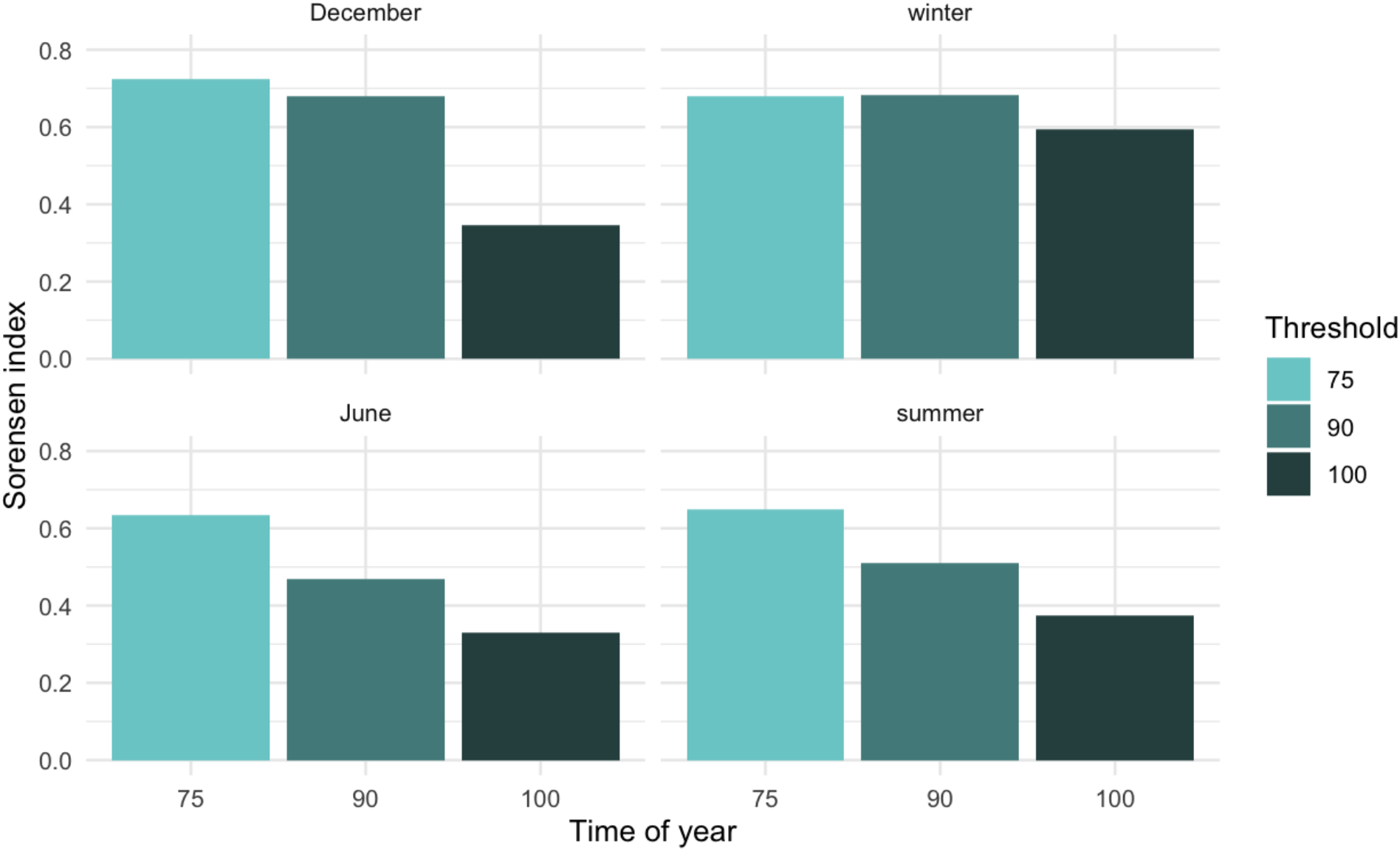
Comparison of the Sørensen index by threshold for one month datasets (December and June) vs two-month datasets (winter and summer)

### 3.4 Statistical analyses for the presence-absence matrices

The 75% thresholds performed the best according to the prediction success for both the summer and the winter dataset. The 90% threshold performed better according to the Sørensen index (Figure 7). The prediction success was higher than the Sørensen index for all thresholds in both datasets. Tables 1 and 2 provide a thorough breakdown of these statistics for each threshold and for both datasets.

**Table 1:** Statistical analysis of presence-absence matrices for Madre Selva winter models

| Threshold | True positive | False positive | True negative | False negative | Prediction success | Sørensen index |
| --- | --- | --- | --- | --- | --- | --- |
| 100 | 106 | 140 | 401 | 4 | 0.779 | 0.595 |
| 90 | 80 | 44 | 497 | 30 | 0.886 | 0.684 |
| 75 | 64 | 14 | 527 | 46 | 0.908 | 0.681 |

**Table 2:**
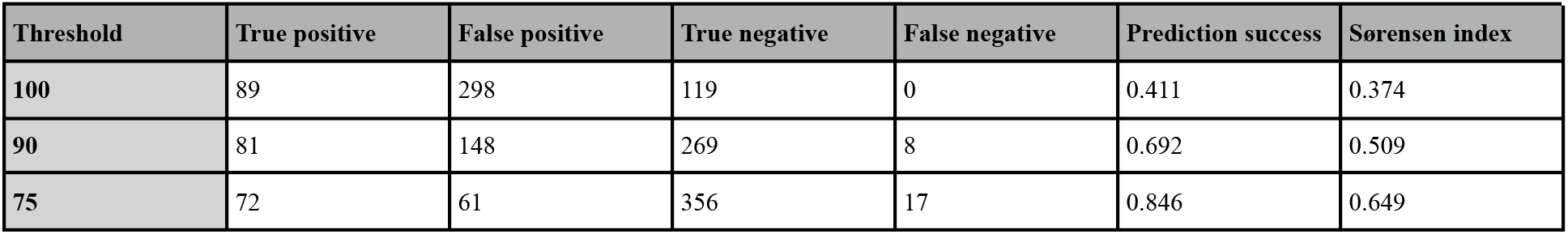
Statistical analysis of presence-absence matrices for San José summer models

| Threshold | True positive | False positive | True negative | False negative | Prediction success | Sørensen index |
| --- | --- | --- | --- | --- | --- | --- |
| 100 | 89 | 298 | 119 | 0 | 0.411 | 0.374 |
| 90 | 81 | 148 | 269 | 8 | 0.692 | 0.509 |
| 75 | 72 | 61 | 356 | 17 | 0.846 | 0.649 |

**Figure 7:**
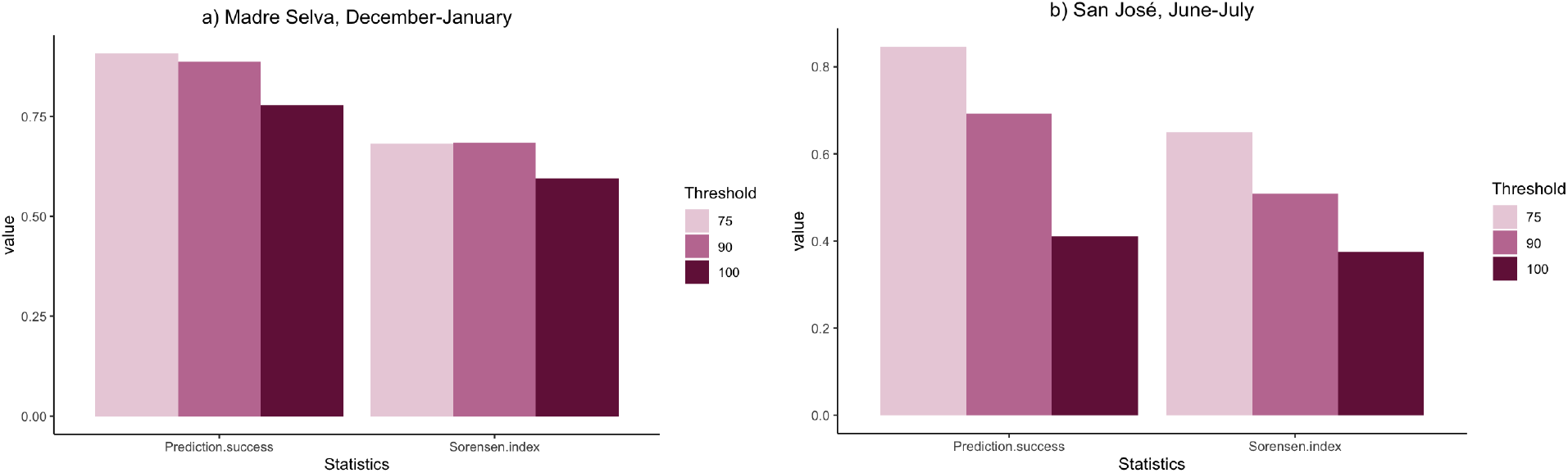
Graphical representation of the statistics for the presence-absence matrices for a) Madre Selva winter models and b) San José summer models

### 3.5 Binning

The amount of data that performed the best was different for the winter and summer dataset (Figure 8). The bins with an intermediate number of observations performed the best for the winter dataset; *i*.*e*., the species with 92-127 observations had the highest prediction success and Sørensen index (Table 3). However, the summer dataset showed an increase in accuracy with more observations; the bins of species with 128-225 had the highest Sørensen index and the bin with 10-29 observations had the highest prediction success (Table 4). The prediction success was high for all bins.

**Table 3:**
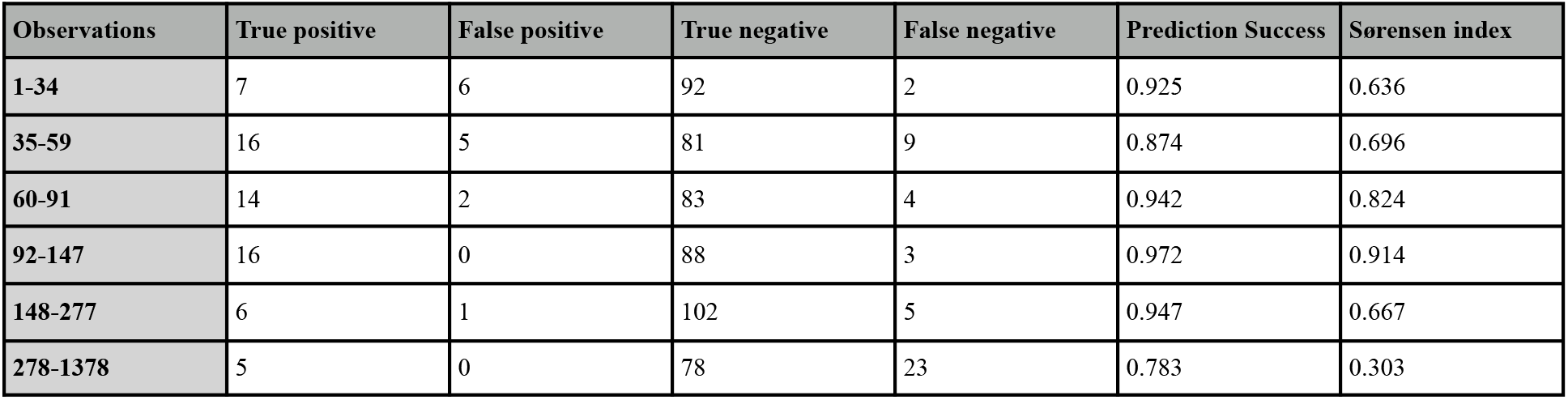
Performance by bins of number of observations for the Madre Selva winter dataset

| Observations | True positive | False positive | True negative | False negative | Prediction Success | Sørensen index |
| --- | --- | --- | --- | --- | --- | --- |
| 1-34 | 7 | 6 | 92 | 2 | 0.925 | 0.636 |
| 35-59 | 16 | 5 | 81 | 9 | 0.874 | 0.696 |
| 60-91 | 14 | 2 | 83 | 4 | 0.942 | 0.824 |
| 92-147 | 16 | 0 | 88 | 3 | 0.972 | 0.914 |
| 148-277 | 6 | 1 | 102 | 5 | 0.947 | 0.667 |
| 278-1378 | 5 | 0 | 78 | 23 | 0.783 | 0.303 |

**Table 4:**
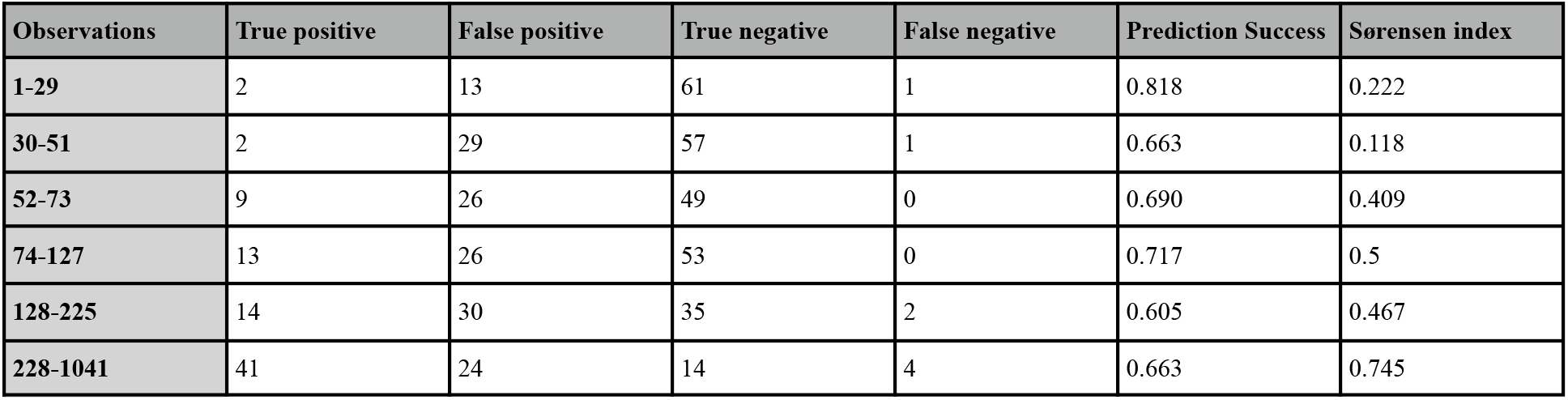
Performance by bins of number of observations for the San José summer dataset

| Observations | True positive | False positive | True negative | False negative | Prediction Success | Sørensen index |
| --- | --- | --- | --- | --- | --- | --- |
| 1-29 | 2 | 13 | 61 | 1 | 0.818 | 0.222 |
| 30-51 | 2 | 29 | 57 | 1 | 0.663 | 0.118 |
| 52-73 | 9 | 26 | 49 | 0 | 0.690 | 0.409 |
| 74-127 | 13 | 26 | 53 | 0 | 0.717 | 0.5 |
| 128-225 | 14 | 30 | 35 | 2 | 0.605 | 0.467 |
| 228-1041 | 41 | 24 | 14 | 4 | 0.663 | 0.745 |

**Figure 8:**
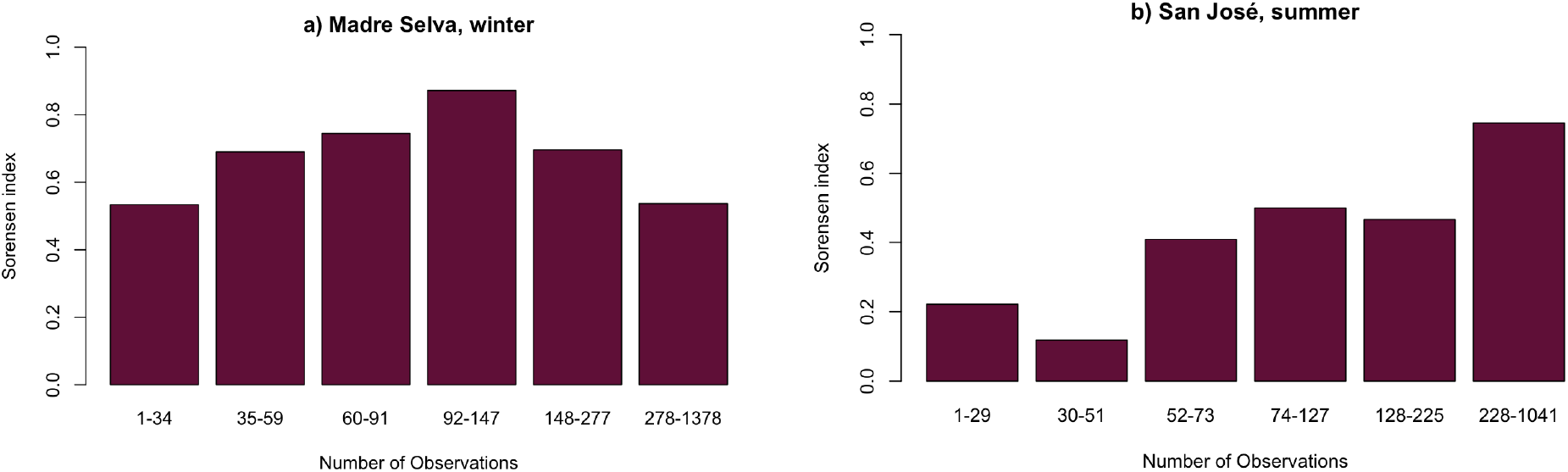
Sørensen index by bins of observations for a) Madre Selva in the winter and b) San José in the summer

## 4. Discussion

My research is one of the first studies to test the predictive abilities of spatial models developed from community science data relative to traditional on-site survey data collected by researchers. I have demonstrated how well community science data, which is inherently patchy, predicts species-level community composition compared to surveys at under-surveyed localities.

I have also evaluated how models performed at different scales in terms of quantity of data, which is an important consideration for the parameters of SDMs. There are a number of studies that examine the decreasing quality of models when data points are scarce (Cayuela et al., 2009; Ruete et al., 2020; Wisz et al., 2008), but this project is one of the first to identify the ideal number of datapoints for stacked SDMs using a presence-only method.

Overall, the SDMs derived from community science data can be good predictors of the community composition at a given locality, given the high prediction success for certain thresholds (up to 90%). However, the success of the models is highly variable across time periods and localities, and certain factors can increase the predictive ability of the models.

I found using field data from a two-month period rather than a one-month period was a more accurate portrayal of the community composition and was more in line with the predictions from the models (Figure 6). This is an indication that we did not identify all of the species that were present and is reflective of real drawbacks of survey methodologies and the inability of researchers to survey an entire community in short time frames. The improvement with increased data from each locality indicates that the models are better at predicting the species than our own short-term field surveys, which can be an important consideration for the data collection process for future projects. Field work can be costly in terms of funds and time, and if species distribution models are more effective at determining community assemblage than researchers themselves, we might see a shift toward primarily using community science data.

The presence-based minimum-volume ellipsoid models are fairly robust to bias, but they are not entirely resistant to biases in the community science data. Certain areas are more poorly surveyed than others because of their accessibility and the higher thresholded models are more biased toward extensively surveyed regions. There are also certain species that are rarely observed, including some that did not even have enough observations to be modeled. In general, rare birds, silent birds, and nocturnal birds tend to be overlooked by observers and are therefore less likely to be detected (Parker, 1991) or, therefore, modeled correctly. However, the binning tests show too much data can also be detrimental. When a site is over-surveyed, it becomes more likely that irregular species or vagrants are included in the composition (Cooper & Soberón, 2018) and some such sought-after species can be over-reported by birders, especially when they are rare (Laney et al., 2021).

In addition to the spatial biases, we can note a large amount of temporal variability across the year. My models for winter and summer offer a good comparison of the distribution of data for the two time periods. The number of observations increases in the winter and decreases in the summer (Figure 2), and this lack of data for the summer time could be a reason why the summer models performed less well than the winter models. One of the main reasons for the differences in data is the high proportion of tourism occurring in the dry season winter months rather than the rainy season summer months (Sistema Nacional de Áreas de Conservación, 2019). Although this is evolving over time, a significant portion of current eBird data for Costa Rica has been collected by tourists, and there is lower tourism during the summer. This implies that summer communities are less well known and more field data needs to be collected in the summer months to make more accurate community models.

It is important to note that the two datasets also differ in their locality, which could have an impact on the quality of the data. Montane communities are easier to model because the environmental conditions can change rapidly with elevation, spatially constraining niches. Our Madre Selva surveys covered an elevation range of ~350 m in the mountains; on the other hand, the San José site is in a dry central valley. The San José area is also heavily urbanized and includes a more disturbed habitat; urbanization reduces species richness, which impacts the bird communities that are present in the San José area (Chace & Walsh, 2006).

There are ways of improving the quality of the models. Using a training area specific to each species would increase the accuracy of their predicted range. Additionally, I chose to use the minimum-volume ellipsoid method to make my models, but for studies using fewer or more reliable data points, there are also other ecological niche modeling techniques that could be used such as Maxent or Generalized Linear Models (Brotons et al., 2004; Pearson et al., 2007; Peterson & Papes, 2006). Furthermore, randomly reducing points to fit within the range of ideal number of observations (100-150) can improve the quality of models as seen by the binning test (Figure 8).

While I focus on community composition, species distribution modeling can also be used to look at community dynamics to better understand their fluctuations over time and the movements of species across various communities. On a practical level, models like these can be used to inform data collection in the field, providing researchers with a better idea of the community composition from the outset. On a larger scale, having an idea of the structure of communities in given areas can be key in determining which areas should be prioritized for protection. The models can predict which regions have high diversity and a concentration of rare or endangered species and can accurately inform current and future conservation efforts.

## 5. Conclusion

By modeling bird species in Costa Rica and Panama and comparing the models’ predictions to *de novo* field observations, I determined that community science data (specifically eBird data) can be a good alternative to data collected by individual observers when certain biases are accounted for. However, community science data, particularly when collected on a worldwide scale, can yield too much data for some species and bias community estimates. My study shows that community science data can be used to make accurate models predicting community composition and confirms the utility of community-science datasets for niche modeling. These analyses can be used for conservation efforts that require an understanding of bird communities in localized and inaccessible areas that may be difficult to reach for extended fieldwork.

## 7. Acknowledgements

I would like to thank John Bates for being an immense help and a wonderful advisor, and the Hackett-Bates Lab at the Field Museum for being more supportive and welcoming than I could have hoped for. I also thank Chespi Elizondo and the Costa Rica Bird Observatories for granting us access to Madre Selva and the resources to gather field data, and the CRBO volunteers for helping with our surveys and contributing independent data. Thanks also to Cathy Pfister for her guidance and enthusiasm throughout the whole process, and Jorge Soberón for providing both his expertise and his code. I am also grateful for everyone in the “Motmots” discussion group for teaching me all I know about research and sometimes making me forget I’m still just an undergrad. I’d like to specifically acknowledge Sara Velásquez Restrepo for helping me through coding for the project and Brian R. Tsuru for letting me distract him from his work at the museum. Finally I would like to thank Holly Garrod whose energy, enthusiasm, and positivity are a constant inspiration, and of course Jacob Cooper, who could not have been a better mentor, and to whom I dedicate all of my accomplishments.

## Footnotes

5 Also referred to as ‘citizen science’

6 University of Chicago & Field Museum of Natural History

7 Costa Rica Bird Observatories & Villanova University

