## Appendix 1 for "Testing the accuracy of species distribution models based on community science data"

Mélusine Velde

#### Species distribution modeling to test the reliability of community science data: code

The goal of this project is to make species distribution models for species in Costa Rica and Panama, and compare the predictions from the models with data that I have collected to test their reliability.

When I was in Costa Rica last December, we performed audio-visual surveys to identify all of the birds in the area we were in. We mostly birded around a field station in Madre Selva, in the San José province. The research station was owned by the Costa Rica Bird Observatories, and was located on private land that is not heavily birded.

#### Section 1: Preparing and cleaning the data

```
library(knitr)
library(lubridate)
library(gridExtra)
library(tidyverse)
library(data.table)
library(fossil)
library(geosphere)
library(rgdal)
select=dplyr::select
```

**1.1 Import/Prepare the files** I have a large file that I acquired from ebird which includes all of the bird observations in Costa Rica and Panama. I reduced the dataset to only include the scientific name, the longitude, latitude, and observation date. Since I am only modeling these species for the month of December (when I was in Costa Rica), I further reduced the data to only include observations from December. From there I further reduced to remove the observations dates.

```
#only include observations from december
cr.pm_december=cr.pm_reduced%>%filter(OBSERVATION.DATE%like%"-12-")
#reduce to scientific name, longitude, latitude
cr.pm_subset = cr.pm_december[,c(1,2,3)]
```

I then want to identify the species that have been found in Madre Selva. This will be my species list for the project. The training area is a shapefile (made in QGIS) which encompasses the accessible geographical area around Madre Selva. Creating a training area allows me to only model the species that are physically able to access Madre Selva. Due to biogeographical barriers such as rivers, mountains, valleys, etc., some species do not have access to Madre Selva; I am eliminating those from my species list because I know they will not be found there.

```
# Identify points within range of training
cr.pm_subsubset=cr.pm_subset%>%
```

```
  select(SCIENTIFIC.NAME,LONGITUDE,LATITUDE)%>%unique()
# read shapefile
EME=readOGR(paste0(dsn, "trainingarea.shp"))

# read file
x.data=cr.pm_subsubset
crs.x=crs(EME)

x.data2=SpatialPoints(x.data[,2:3],proj4string = crs.x)

# is it inside the polygon?
inside.m=!is.na(over(x.data2,as(EME,"SpatialPolygons")))

# create new dataframe for the training area species
ta.species=x.data[which(inside.m==T),]

sp=ta.species
#write_csv(sp,paste0(filepath, "species.trainingarea", ".csv"))
```

I make a species list from the list of birds found in the training area.

```
allspecieslist=sp%>%
  select(SCIENTIFIC.NAME)%>%unique()
#write.csv(allspecieslist,paste0(filepath,"/all.uniquespecies",".csv"))
```

```
all.specieslist=read_csv(paste0(filepath, "all.uniquespecies.csv"))
all.specieslist=all.specieslist[c(2)]
all.specieslist=all.specieslist$SCIENTIFIC.NAME

all.specieslist=as.character(all.specieslist)
print(all.specieslist[509:529])
```

```
##  [1] "Oncostoma cinereigulare"
##  [2] "Empidonax sp."
##  [3] "Aphanotriccus capitalis"
##  [4] "Dendrocolaptinae sp."
##  [5] "Piranga ludoviciana"
##  [6] "Setophaga townsendi"
##  [7] "Dendrocygna autumnalis"
##  [8] "Vireo solitarius"
##  [9] "Zentrygon chiriquensis"
## [10] "Patagioenas speciosa"
## [11] "Phalaropus tricolor"
## [12] "Columba livia"
## [13] "Falco sp."
## [14] "Morococcyx erythropygus"
## [15] "Porphyrio martinica"
## [16] "Thalasseus sandvicensis"
## [17] "Contopus virens"
## [18] "Lampornis calolaemus/castaneoventris"
## [19] "Phylloscartes superciliaris"
## [20] "Baryphthengus martii"
## [21] "Psittaciformes sp. (parakeet sp.)"
```

**1.2 Make legible species list** I then sort the data by species and create a csv file of all of the species I will be modeling. A number of observations have to be removed because the species names are either not complete or are hybirds (which I will not be modeling).

```
#REMOVE ALL SPECIES WITH ADDITIONAL CHARACTERS/TERMS

for(i in 1:length(all.specieslist)){
  genus=strsplit(all.specieslist[i]," ")[[1]][1]
  species=strsplit(all.specieslist[i]," ")[[1]][2]
  filename=paste0(genus,"-",species,".csv")

  x=cr.pm_subsubset%>%filter(SCIENTIFIC.NAME==all.specieslist[i])
  #write.csv(x,paste0(SaveDir,filename))
}
```

**1.3 Rarify** We want only unique combinations of species/longitude/latitude so as not to overfit our models. If the same species of bird has been recorded many times in the same spot, the model will see that area as much more suitable than others. Instead we want all of the locations where each species has been found.

```
melusine.distance=function(input,distance.km){
  input=input[order(input[,2]),]
  input2=input
  for(i in 1:nrow(input)){
    #if the i is greater than columns break loop
    if(nrow(input)<i){next}

    # create a matrix of distances for all points
    y=as.numeric(as.matrix(earth.dist(as.data.frame(input)))[,i])
    # find rows less than distance
    index=as.numeric(which(y<distance.km&y>0)) #remove identity cell!
    # print row reduction
    # reduce data frame
    # run again
    if(length(index)>0){
      print(paste0(nrow(input)," rows reduced by ",length(index)))
      input=input[-index,]
  }
  }
  plot(input2$LONGITUDE,input2$LATITUDE,pch=".",col="black",asp=1,xlab="Longitude",ylab="Latitude")
  points(input$LONGITUDE,input$LATITUDE,pch=20,col="red")

  return(input)
}
```

```
for(i in 1:length(file.names)){
  genus=strsplit(file.names[i],"-")[[1]][1]
  epithet=strsplit(file.names[i],"-")[[1]][2]
  species=strsplit(epithet,"[.]")[[1]][1]
  filename=paste0(genus,"-",species,"_rarified.csv")
  df=read_csv(paste0(path,file.names[i]))
  df.red=df%>%select(LONGITUDE,LATITUDE)%>%as.data.frame()
  x=melusine.distance(input= df.red, distance.km = 1)
  #write.csv(x,paste0(SaveDir,filename),row.names = F)
}
```

**1.4 Create artificial absence** I removed all points within 10 km of the Madre Selva research station (-83.867218, 9.661011). This is important because I do not want to use my own data (which is already in eBird) to make the models; I want to be able to compare what is predicted to be there based on observations from the rest of the country.

```
p1=as.vector(c((-83.9837813), (9.9554869)))

artificial.absence=function(input,distance.km){
  input=as.matrix(input)
  x.rows=NULL
  for(i in 1:nrow(input)){
    p2=c(input[i,1], input[i,2])
    d.x=distMeeus(p1,p2)/1000
    if(d.x<distance.km){
      if(is.null(x.rows)){x.rows=i}else{
        x.rows=c(x.rows,i)
      }
    }else(next)
  }
  if(is.null(x.rows)){
    print("No rows removed.")
    return(input)
  }else{
    print(paste0(length(x.rows)," rows removed."))
    input=input[-x.rows,]
    return(input)
  }
}

for(i in 1:length(file.names.rarified)){
  genus=strsplit(file.names.rarified[i],"-")[[1]][1]
  epithet=strsplit(file.names.rarified[i],"-")[[1]][2]
  species=strsplit(epithet,"[.]")[[1]][1]
  filename=paste0(genus,"-",species,"_absence.csv")
  df=read.csv(paste0(path,file.names.rarified[i]))
  x=artificial.absence(input= df, distance.km = 10)
  #write.csv(x,paste0(SaveDir,filename),row.names = F)
}
```

### Section 2: Preparing data layers (NDVI and envirem)

NDVI layer packages

```
library("data.table")
library("gdalUtils")
library("rgdal")
library("raster")
library("nloptr")
library("processx")
library("fs")
library("MODIS")
library("sf")
library("tibble")
library("envirem")
```

```
library("tidyverse")
library("data.table")
library("rgdal")
#install.packages("gdalUtils")
library("gdalUtils")
#install.packages("RGtk2")
#library("RGtk2")
#install.packages("MODIStsp")
#library("MODIStsp")

#install.packages("~/Downloads/gWidgets_0.0-54.tar.gz/")
#install.packages('gWidgets',repos='http://cran.us.r-project.org')
#install.packages("gWidgetsRGtk2")

#library(devtools)
#install_url('https://cran.r-project.org/src/contrib/Archive/velox/velox_0.2.0.tar.gz')
library("velox")

#library(devtools)
#install_url('https://cran.r-project.org/src/contrib/Archive/gWidgets/gWidgets_0.0-54.tar.gz')
#library("gWidgets")
#library(devtools)
#install_url('https://cran.r-project.org/src/contrib/Archive/gWidgetsRGtk2/gWidgetsRGtk2_0.0-86.tar.gz'
#library("gWidgetsRGtk2")
```

**2.1 Identifying tiles for study area** First, I had to create an account with earthdata to have access to the tiles
Then I used the USGS earthdata website (urs.earthdata.nasa.gov) to identify the necessary tiles that cover my entire stufy area (Costa Rica and Panama); they make it very easy, so you can draw a study area and they identify the tiles.
I used MOD13A2 (MODIS/Terra Vegetation Indices 16-Day L3 Global 1km SIN Grid V006); these are environmental datalayers showing NDVI (landcover). There are 3 tiles corresponding to CR and PN

```
#create a study area as a shapefile in QGIS, then designate it as the study area
studyarea=read_sf(paste0(filepath2, "study_area2.shp"))
plot(studyarea)
```

## id

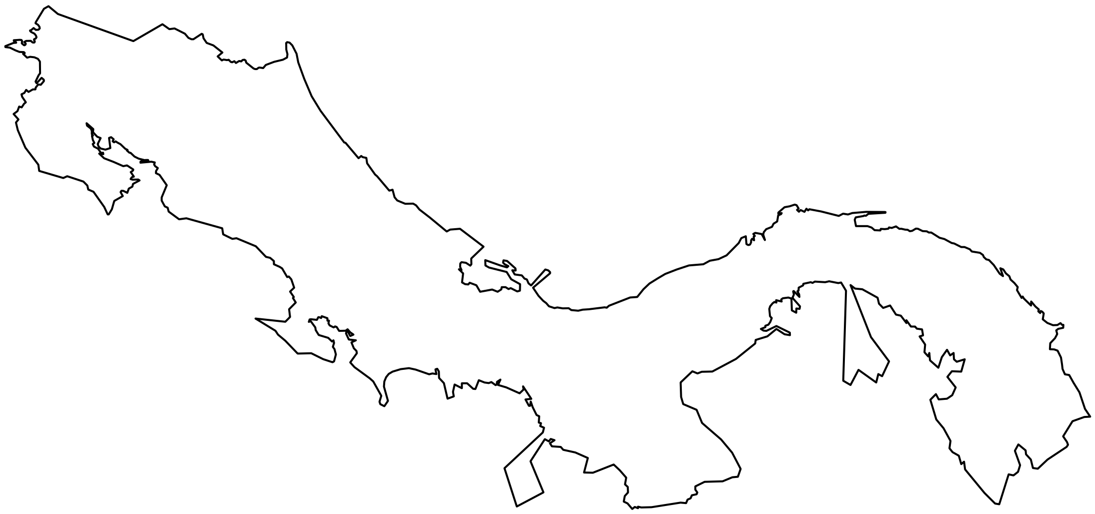

```
#MODIS::EarthdataLogin(usr="melusinevelde",pwd="Earthdata23")
gdal_chooseInstallation(hasDrivers="HDF4")
```

```
## [1] 1
```

```
#convert bounding box to modis coordinates
zoom2=studyarea %>% st_transform(crs=paste("+proj=sinu +lon_0=0 +x_0=0 +y_0=0",
                              "+a=6371007.181 +b=6271007.181 +units=m +no_defs"))
#get tile names
tiles=getTile(zoom2)
#print tile names
tiles@tile
```

```
## [1] "h09v08" "h10v08" "h09v07"
```

**2.2 Convert .hdf to .tif** Once I have the data for NDVI for each tile, I convert the .hdf files to .tif (This chunk did not work on my computer, I had to do it on Jacob's)

```
# run on Jacob's Computer
hdf_files=list.files(paste0(GIS,"NDVI/hdf/"),pattern="*.hdf")

for(i in 1:length(hdf_files)){
  # get subdata layers
  print(paste0("Working:",(round(i/length(hdf_files),2)*100),"%"))
  sds=get_subdatasets(hdf_files[i])

  # find NDVI layer
  k=which(sds %like% "NDVI")

  # get coverage date
  x.date=strsplit(sds[k],"[.]")[[1]][2]
  x.year=strsplit(x.date,"")[[1]][2:5]
  x.year=paste0(x.year,collapse="")
```

```
  x.day=strsplit(x.date,"")[[1]][6:8]
  x.day=paste0(x.day,collapse="")
  # get tile location
  x.pos=strsplit(sds[k],"[.]")[[1]][3]

  new.filename=paste0(GIS,"NDVI/tif/","NDVI_modis_",x.year,"-",x.day,"_",
                      x.pos,"_gridded.tif")

  hdf.x=readGDAL(sds[k],options=c("RASTERXDIM=4","RASTERYDIM=3","RASTERBDIM=2","RASTER4DIM=1","RASTER5DI

  r=raster(hdf.x)

  #writeRaster(x=r,filename=paste0(new.filename),overwrite=T)
}
```

**2.3 Combine tifs into per-date set of 3** Each date has 3 tiles (correponding to different parts of Costa Rica and Panama). I combined the three tiles into one large tile to get one per date.

```
geotiff=list.files(paste0(GIS,"NDVI/tif/gridded/"),pattern="*_gridded.tif")

# first, to merge by date
# get unique dates

for(i in 1:length(geotiff)){
  date=strsplit(geotiff[i],"_")[[1]][3]
  if(i==1){dates=date}else{dates=c(dates,date)}
  dates=unique(dates)
}

# show NDVI date slices
dates
```

```
##  [1] "2003-337" "2003-353" "2004-337" "2004-353" "2005-337" "2005-353"
##  [7] "2006-337" "2006-353" "2007-337" "2007-353" "2008-337" "2008-353"
## [13] "2009-337" "2009-353" "2010-337" "2010-353" "2011-337" "2011-353"
## [19] "2012-337" "2012-353" "2013-337" "2013-353" "2014-337" "2014-353"
## [25] "2015-337" "2015-353" "2016-337" "2016-353" "2017-337" "2017-353"
## [31] "2018-337" "2018-353" "2019-337" "2019-353"
```

```
setwd(paste0(GIS,"NDVI/tif/gridded/"))

for(i in 1:length(dates)){
  date=dates[i]

  index=which(geotiff %like% date)
  if(length(index)<3){next}

  # stitch together

  x.geo=geotiff[index]

  #mosaic_rasters(gdalfile = x.geo,of="GTiff",
                #dst_dataset = paste0("NDVI_",date,"_combined.tif"))
}
```

**2.4 Stacking tiles by year** Get dates corresponding to each year

```
geotiff=list.files(paste0(GIS, "NDVI/tif/combined/"),pattern="*_combined.tif")
for(i in 1:length(geotiff)){
  date=strsplit(geotiff[i],"_")[[1]][2]
  year=strsplit(date,"-")[[1]][1]
  if(i==1){dates=year}else{dates=c(dates,year)}
  dates=unique(dates)
}

# show NDVI date slices
dates
```

**2.5 Get maximum and minimum NDVI for each date** Stack the years together to get a mininum NDVI and a maximum NDVI for each year. December is at the cusp between the dry season and the rainy season, so it's better to make a maximum and minimum layer to account for the large potential variation. Using both ensures that we capture the extrema that the birds can handle

```
x.gtif=paste0(GIS,"NDVI/tif/combined/",geotiff)

for(i in 2:length(dates)){
  date=dates[i]

  # which files contain that year?
  index=which(x.gtif %like% date)

  # get full filenames
  x.geo=x.gtif[index]

  # stack all raster files for year
  r=raster::stack(x.geo)

  # calculate minimum raster
  r.min=calc(r,fun=min)
  #writeRaster(r.min,paste0(GIS,"NDVI/tif/combined/yearly_min/NDVI_",date,"_yearly_min.tif"), overwrite
  rm(r.min)

  # calculate maximum raster
  r.max=calc(r,fun=max)
  #writeRaster(r.max,paste0(GIS,"NDVI/tif/combined/yearly_max/NDVI_",date,"_yearly_max.tif"), overwrite
  rm(r.max)

  # free memory
  rm(r)

  print(paste0("Year completed: ",date))
}

# remove all files of a type from folder
# open terminal
# navigate to folder
# rm *.grd
```

**2.6 Aligning variable rasters** Create "blank canvas" for reprojecting the environmental variables This is important because all of the variables need to be aligned on the same projection (4326) to match up correctly on the maps

```
e.cr=raster(paste0(GIS,"NDVI/tif/combined/yearly_max/NDVI_2003_yearly_max.tif"))
map_proj=st_crs(4326)
canvas=projectExtent(e.cr,crs = map_proj$proj4string)
test=projectRaster(e.cr,canvas,method="bilinear")
res.x=res(test)
ext.x=extent(test)
plot(test)
```

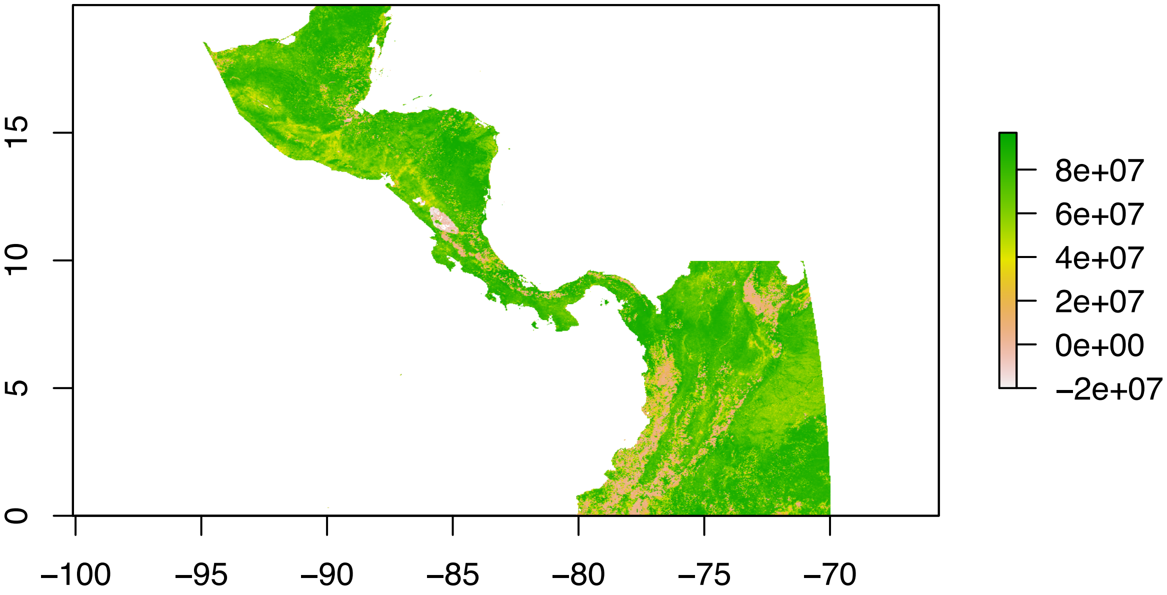


```
rm(e.cr)
rm(test)
```

**2.7 Reproject envirem layers onto this blank canvas** I got 7 datalayers from the Envirem website The layers used are:
- aridityIndexThornthwaite
- continentality
- embergerQ
- maxTempColdest
- minTempWarmest
- PETDriestQuarter
- PETWettestQuarter

There are a lot of other variables, but some of them are heavily correlated using all of them would cause the models to be biased toward those. These ones explain the most amount of variation so these ones were used to keep it simple.

```
env_30=paste0(GIS,"env_variables/", list.files(paste0(GIS, "env_variables/")),,pattern="*_combined.tif"

f1=which(env_30 %like% "aridityIndexThornthwaite.tif")
f2=which(env_30 %like% "continentality.tif")
f3=which(env_30 %like% "embergerQ.tif")
f4=which(env_30 %like% "maxTempColdest.tif")
f5=which(env_30 %like% "minTempWarmest.tif")
```

```
f6=which(env_30 %like% "PETDriestQuarter.tif")
f7=which(env_30 %like% "PETWettestQuarter.tif")

files=env_30[c(f1,f2,f3,f4,f5,f6,f7)]

for(i in 1:length(files)){
  bio=raster(files[i])
  name=strsplit(files[i],"_")[[1]][6]
  x3=projectRaster(from=bio,to=canvas,method="bilinear",
                   res=res.x,crs=map_proj$proj4string)
  rm(bio)
  #writeRaster(x3,paste0(GIS,"env_variables/reprojected/",name,".tif"), overwrite=T)
}
```

```
ndvi_min=paste0(GIS,"NDVI/tif/combined/yearly_min/",list.files(paste0(GIS,"NDVI/tif/combined/yearly_min
ndvi_max=paste0(GIS,"NDVI/tif/combined/yearly_max/",list.files(paste0(GIS,"NDVI/tif/combined/yearly_max

for(i in 1:length(ndvi_min)){
  bio=raster(ndvi_min[i])
  nam=strsplit(ndvi_min[i],"/")[[1]][12]
  name=strsplit(nam,"[.]")[[1]][1]
  x3=projectRaster(from=bio,to=canvas,crs=map_proj$proj4string,res=res.x,method="bilinear")
  #writeRaster(x3,paste0(GIS,"NDVI/tif/combined/yearly_min/reprojected/",name, "_reprojected", ".tif"))
}

for(i in 1:length(ndvi_max)){
  bio=raster(ndvi_max[i])
  nam=strsplit(ndvi_max[i],"/")[[1]][12]
  name=strsplit(nam,"[.]")[[1]][1]
  x3=projectRaster(from=bio,to=canvas,crs=map_proj$proj4string,res=res.x,method="bilinear")
  #writeRaster(x3,paste0(GIS,"NDVI/tif/combined/yearly_max/reprojected/",name, "_reprojected", ".tif"))
}

min_files=paste0(GIS,"NDVI/tif/combined/yearly_min/reprojected",list.files(paste0(GIS,"NDVI/tif/combine
max_files=paste0(GIS,"NDVI/tif/combined/yearly_max/reprojected",list.files(paste0(GIS,"NDVI/tif/combine
```

##### 2.8 Convert NDVI to present projection

```
landmin.av=calc(landcover.min,fun=mean)
#writeRaster(landmin.av,paste0(GIS,"env_variables/", "NDVImin_average", ".tif"))
landmax.av=calc(landcover.max,fun=mean)
#writeRaster(landmax.av,paste0(GIS,"env_variables/", "NDVImax_average", ".tif"))
```

##### 2.9 average all max and all min NDVI across all years; in addition to the 7 envirem datalayers, these are part of the environmental datalayers

```
files=paste0(GIS,"env_variables/",list.files(paste0(GIS,"env_variables/"),pattern="*.tif"))
for(i in 1:length(files)){
  raster.files=raster(files[i])
  nam=strsplit(files[i],"/")[[1]][9]
```

```
  name=strsplit(nam, "[.]")[[1]][1]
  x=crop(raster.files, studyarea)
  #writeRaster(x, paste0(GIS, "env_variables/test/", name, "_cropped", ".tif"), overwrite=T)
}
```

##### 2.10 clip all layers to CR and PM study area

**2.11 Extract environmental point characters** 1 deg is about 110 km according to the NOAA website This step creates a dataframe of all the environmental conditions (based on the 9 datalayers) for each locality (latitude, longitude).

```
# NDVI landcover files

landcover.min=paste0(GIS,"NDVI/tif/combined/yearly_min/reprojected/",list.files(paste0(GIS,"NDVI/tif/co
landcover.max=paste0(GIS,"NDVI/tif/combined/yearly_max/reprojected/",list.files(paste0(GIS,"NDVI/tif/co

landcover.min=stack(landcover.min)

landcover.min=names(landcover.min) %>%
  str_extract("([0-9]+)") %>%
  paste0("y",.) %>%
  setNames(landcover.min, .)

landcover.max=stack(landcover.max)

landcover.max=names(landcover.max) %>%
  str_extract("([0-9]+)") %>%
  paste0("y",.) %>%
  setNames(landcover.max, .)

# ENVIREM files
envirem.files=paste0(GIS,"env_variables/env_reprojected/",list.files(paste0(GIS,"env_variables/env_repr

envirem=stack(envirem.files)
```

```
print(envirem)
```

Packages

```
library(dggridR)
library(ranger)
library(scam)
library(PresenceAbsence)
library(data.table)
library(verification)
library(maptools)
library(ebirdst)
library(fields)
library(tidyverse)
library(sf)
library(raster)

select=dplyr::select
projection=raster::projection
map=purrr::map
```

More packages

```
library(tidyverse)
library(raster)
library(MASS)
library(sf)
library(velox)
library(gdalUtils)
```

**2.12 Match point data with environmental conditions** This is extracting the environmental conditions at each observation

```
x.sample=raster(paste0(GIS,"NDVI/tif/combined/yearly_min/reprojected/NDVI_2018_yearly_min_reprojected.ti
#define neighborhood size - 3 km multiplied by resolution
# distance radius around each point
dist_r=2/110

# number of cells around each point
cell_r=ceiling((2/110)/max(res(x.sample)))
```

It is important to first remove all the files that have fewer than 1 + the number of environmental variables

```
occs=paste0(filepath,"artificial.absence/",list.files(paste0(filepath,"artificial.absence/"),pattern="*
occs=unlist(occs)

current=paste0(GIS,"env_variables/",list.files(paste0(GIS,"env_variables/"),pattern="*.tif"))
current=stack(current)

for(i in 1:length(occs)){

  ebird=read.csv(occs[i])

  #add a checklist_id
  v.x=paste0('x',1:nrow(ebird))
  ebird$checklist_id=v.x

  # we do not need year-by-year
  # conditions are across period

  ebird_buff_noyear=ebird %>%
    distinct(checklist_id,LATITUDE,LONGITUDE) %>%
    st_as_sf(coords=c("LONGITUDE","LATITUDE"),crs=4326) %>%
    st_buffer(dist=dist_r)

  locs=st_set_geometry(ebird_buff_noyear,NULL) %>%
    mutate(id=row_number())

  envirem_checklists=velox(current)$extract(ebird_buff_noyear,df=T) %>%
    set_names(c("id","aridityIndexThornthwaite_reprojected_cropped",
                "continentality_reprojected_cropped","embergerQ_reprojected_cropped",
                "maxTempColdest_reprojected_cropped","minTempWarmest_reprojected_cropped",
                "NDVImax_average_cropped","NDVImin_average_cropped",
                "PETDriestQuarter_reprojected_cropped","PETWettestQuarter_reprojected_cropped")) %>%
    inner_join(locs,.,by="id")
```

```
  # number of checklists maintained to here

  envirem_checklists=envirem_checklists %>% dplyr::select(-id) %>%
    filter(!is.na(minTempWarmest_reprojected_cropped))

  env.land=aggregate(. ~ checklist_id,envirem_checklists,mean)

  ebird2=inner_join(ebird,env.land,by="checklist_id")

  name1=strsplit(occs[i],"/")[[1]][8]
  name2=strsplit(name1,"_")[[1]][1]
  #write_csv(ebird2,paste0(filepath,"envirem.extracts/",name2,"_clipped-extracts.csv"))
}
```

#### Section 3: Predictive surfaces

Make a single predictive surface where each point (longitude/latitude) has environmental conditions associated with it

```
# create single predictive surface each time slice
studyarea=read_sf(paste0(filepath, "studyarea_allCR-PA/study_area2/study_area2.shp"))
plot(studyarea)
```

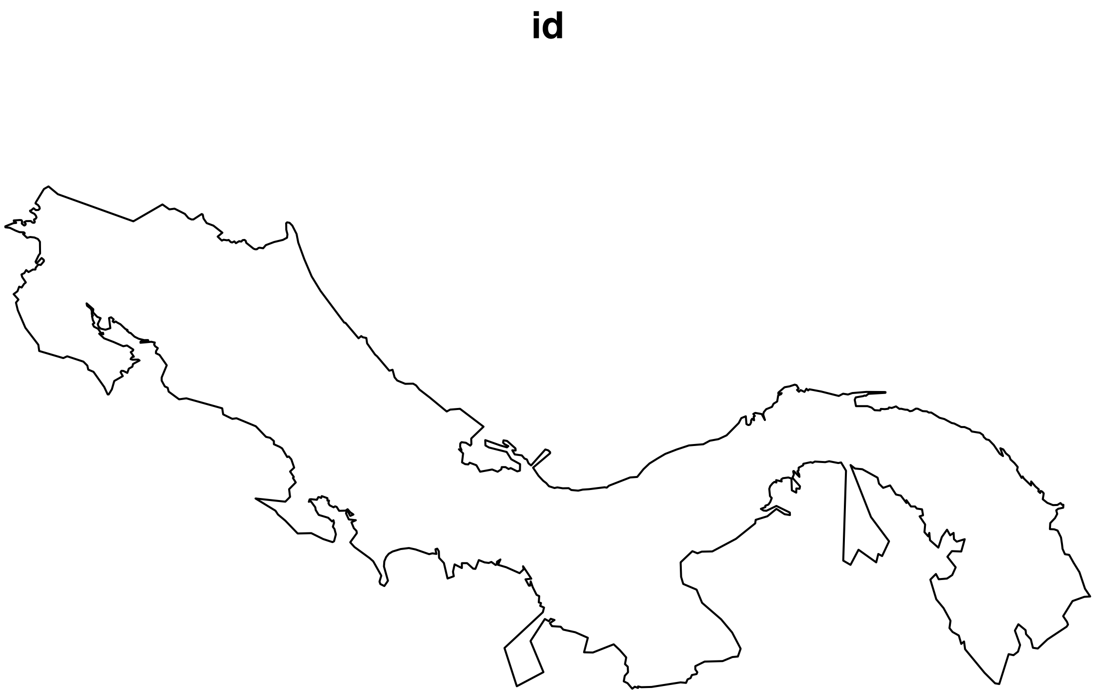


```
current=paste0(GIS,"env_variables/",list.files(paste0(GIS,"env_variables/"),pattern="*.tif"))
current=stack(current)

names=set_names(c("aridityIndexThornthwaite_reprojected_cropped",
                "continentality_reprojected_cropped","embergerQ_reprojected_cropped",
                "maxTempColdest_reprojected_cropped","minTempWarmest_reprojected_cropped",
                "NDVImax_average_cropped","NDVImin_average_cropped",
                "PETDriestQuarter_reprojected_cropped","PETWettestQuarter_reprojected_cropped"))
```

```
test=current[[1]]

r2=studyarea %>%
  rasterize(test) %>%
  trim()

r_centers=rasterToPoints(r2,spatial=T)

r_cells=r_centers %>%
  st_as_sf() %>%
  transmute(id=row_number())

# extract values for points only
lc_vlx=velox(current[[names]])
lc_extract=lc_vlx$extract_points(r_cells)
lc_extract=lc_extract %>%
  as.data.frame() %>%
  set_names(names) %>%
  mutate(id=row_number())

bio_coords=r_cells %>%
  st_coordinates %>%
  as.data.frame() %>%
  mutate(id=row_number()) %>%
  rename(longitude=X,latitude=Y)

bio_coords=inner_join(lc_extract,bio_coords,by="id")

#write_csv(bio_coords,paste0(filepath,"Current_predictive_surface.csv"))
```

#### Section 4: Minimum volume ellipsoids

To estimate the parameters of my models, I used the minimum volume ellipsoid method which is trained with Mahalanobis distances. It creates a multidimensional ellipsoid that encloses all of the data points. The center of the ellipsoid represents the most suitable habitat for the species (based on suitable environmental conditions), and the farther a data point is from the center of the ellipsoid, the less likely it is that the species in question will be found there.

```
library(MASS)
library(ellipse)
library(raster)
library(tidyverse)
library(data.table)
library(maptools)
library(dplyr)
library(rgdal)
```

Code provided by Jacob from Jorge Soberón

```
#MAJA function
maja=function(p,m,s)((p-m)%*%s%*%t(p-m))^0.5

#Quantile function
```

```
##Double check function? divide by 1 is 1...
##changed to 4, for quantiles
NDquantil=function(nD,level){
  return(round(nD*level))
}
```

Designate the species list

```
env.files=paste0(filepath,"envirem.extracts/",list.files(paste0(filepath,"envirem.extracts/"),pattern="
names=NA
for(i in 1:length(env.files)){
  nombre=strsplit(env.files,"_")[[i]][1]
  index=length(strsplit(nombre,"/")[[1]])
  names[i]=strsplit(nombre,"/")[[1]][index]
}

names=unique(names)
```

Create EVIREM modern models

```
red.x=1000000 # reduction factor for NDVI

# cannot solve matrices if values too disparate
pred.x=read_csv(paste0(GIS,"Current_predictive_surface.csv"))
```

```
pred.x$NDVImax_average_cropped=pred.x$NDVImax_average_cropped/red.x
pred.x$NDVImin_average_cropped=pred.x$NDVImin_average_cropped/red.x

nc=nrow(pred.x)

#pred.x=pred.x[order(pred.x$id),]
```

```
setwd(paste0(filepath))
env.files=paste0(filepath,"envirem.extracts/",list.files(paste0(filepath,"envirem.extracts/"),pattern="
output=paste0(filepath,"output_trainingarea/")
```

Remove files that have fewer than 11 species

```
for(i in 1:length(env.files)){
  file=read.csv(env.files[i])
  rows=nrow(file)
  if(nrow(file)<11){
    print(env.files[i])
  }
}
```

Make the species distribution maps with different thresholds. The thresholded models are binary predictions, and the non-thresholded model is a gradient of suitable to unsuitable habitat. Thresholding reflects the confidence levels; for thresholding at 90% for example, we are assuming that only 90% of the data points are accurate. This is a way of removing erroneous data.This accounts for vagrants and misidentifications.

```
pts=pred.x%>%dplyr::select(longitude,latitude)

for(i in 651:length(env.files)){
  # get species name
  name.x=strsplit(env.files[i],"/")[[1]][8]
  name=strsplit(name.x,"_")[[1]][1]
```

```r
# load environmental data

vals=read_csv(env.files[i])

x.coords=vals%>%dplyr::select(LONGITUDE,LATITUDE)
vals=vals%>%dplyr::select(-LONGITUDE,-LATITUDE,-checklist_id)

vals$NDVImin_average_cropped=vals$NDVImin_average_cropped/red.x
vals$NDVImax_average_cropped=vals$NDVImax_average_cropped/red.x

n1=NDquantil(nrow(vals),0.9)

rob=cov.rob(vals,quantile.use=n1)
is.na(vals)

mu2=matrix(rob$center,nrow=1)%>%as.matrix()
s1=rob$cov
invs2=solve(s1)%>%as.matrix()

dT1=matrix(0,ncol=1,nrow=nc)

# load projection raster

valsT=pred.x%>%
  #dplyr::select(-id,-longitude,-latitude)%>% # remove coordinates
  dplyr::select(colnames(vals)) # make the order the same for both

valsT1=as.matrix(valsT)
rm(valsT)

for(j in 1:nrow(valsT1)){
  dT1[j,1]=maja(valsT1[j,],mu2,invs2)
}

pts2=cbind(pts,dT1)
coordinates(pts2)=~longitude+latitude
gridded(pts2)=T
rast.x=rasterFromXYZ(pts2)

M=readOGR(paste0(filepath,"studyarea_allCR-PA/study_area2/study_area2.shp"))

cr1=raster::crop(rast.x,M)
r_pred=raster::mask(cr1,M)

#plot(r_pred,main=name)
#points(x=x.coords$longitude,
     #y=x.coords$latitude,pch=19)

#writeRaster(r_pred,
            #file.path(paste0(output,"all/",name,"_all-pts.tif")), overwrite=T)

# prepare to threshold
rf.vals=raster::extract(x=r_pred,y=x.coords)
```

```
  if(sum(is.na(rf.vals))>0){
    rf.vals=rf.vals[-which(is.na(rf.vals))]}
  rf.vals=rf.vals[order(rf.vals)]

  # thresholding on confidence in the points
  # not based with confidence of furthest point

  n.t=length(rf.vals)

  thresh.100=max(rf.vals)
  thresh.90=max(rf.vals[1:round(0.9*n.t)])
  thresh.75=max(rf.vals[1:round(0.75*n.t)])

  r_pred_100=r_pred
  r_pred_100[r_pred_100<=thresh.100]=1
  r_pred_100[r_pred_100>thresh.100]=0

  # save the raster
 # writeRaster(r_pred_100,file.path(paste0(output,"100/",name,"_all-pts_100.tif")), overwrite=T)

  r_pred_90=r_pred
  r_pred_90[r_pred_90<=thresh.90]=1
  r_pred_90[r_pred_90>thresh.90]=0

  #points(obs.x[,-1],pch=19)

  # save the raster
 # writeRaster(r_pred_90,file.path(paste0(output,"90/",name,"_all-pts_90.tif")),overwrite=T)

  r_pred_75=r_pred
  r_pred_75[r_pred_75<=thresh.75]=1
  r_pred_75[r_pred_75>thresh.75]=0

  #points(obs.x[,-1],pch=19)

  # save the raster
  #writeRaster(r_pred_75,file.path(paste0(output,"75/",name,"_all-pts_75.tif")),overwrite = TRUE)

}
```

There are a lot of models, so here is an example of one of the species (Zonotrichia capensis)
In the first model, we have a gradient of suitable (white) to unsuitable (green). The scale shows how far the point is from the center of the minimum volume ellipsoid; the farther the point, the least likely it is that the habitat is suitable for the species.
The second model is thresholded at 75%. The green means suitable habitat (and that it could theoretically be found there) and grey means it is unsuitable and will not be found there. The third and fourth models are thresholded at 90% and 100%.
It is important to note however, that most of the green areas might not be accessible to the species, and while the habitat might be suitable, the bird might not live in that area. For my project, it does not matter if I project these models onto the entire country or just onto my training area, because I will only be looking at the species predicted to be in Madre Selva, which was inside my training area.

```
#suitability gradient
plot(r_pred)
```

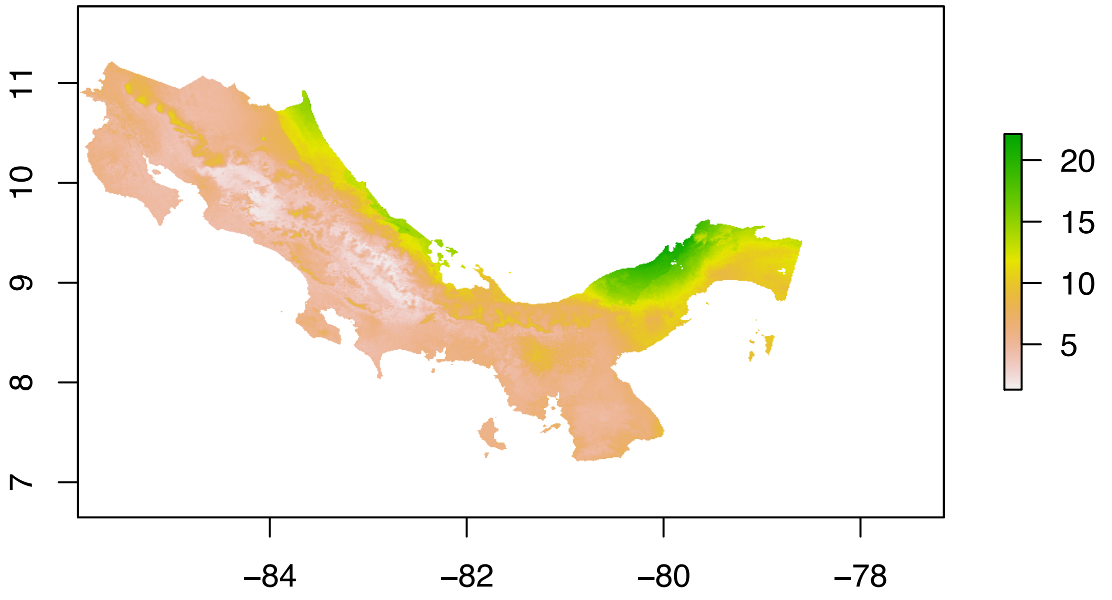


```
#75% threshold
plot(r_pred_75)
```

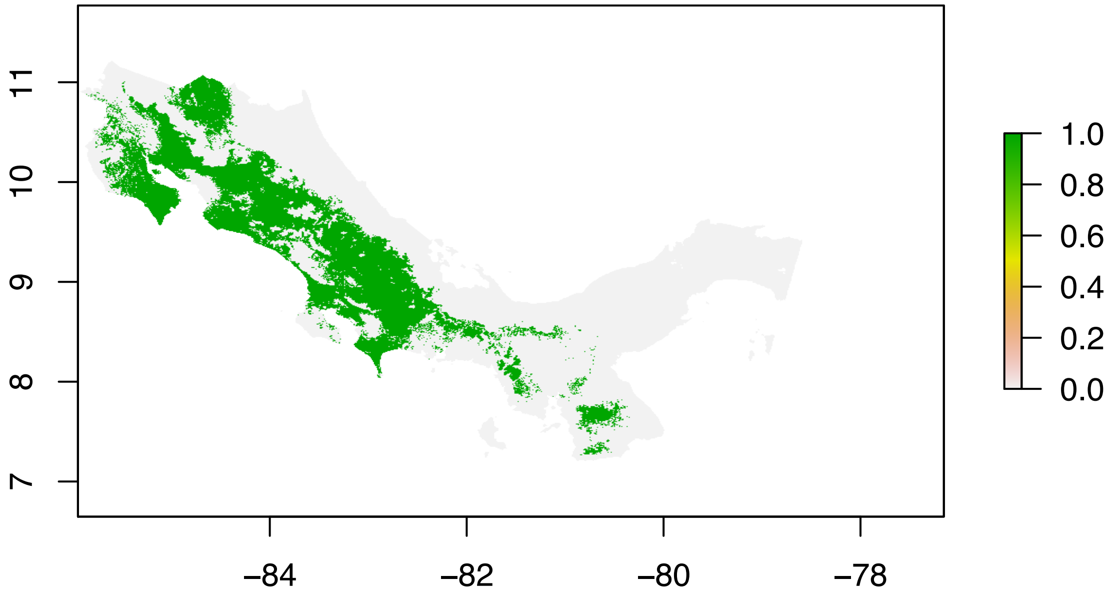


```
#90% threshold
plot(r_pred_90)
```

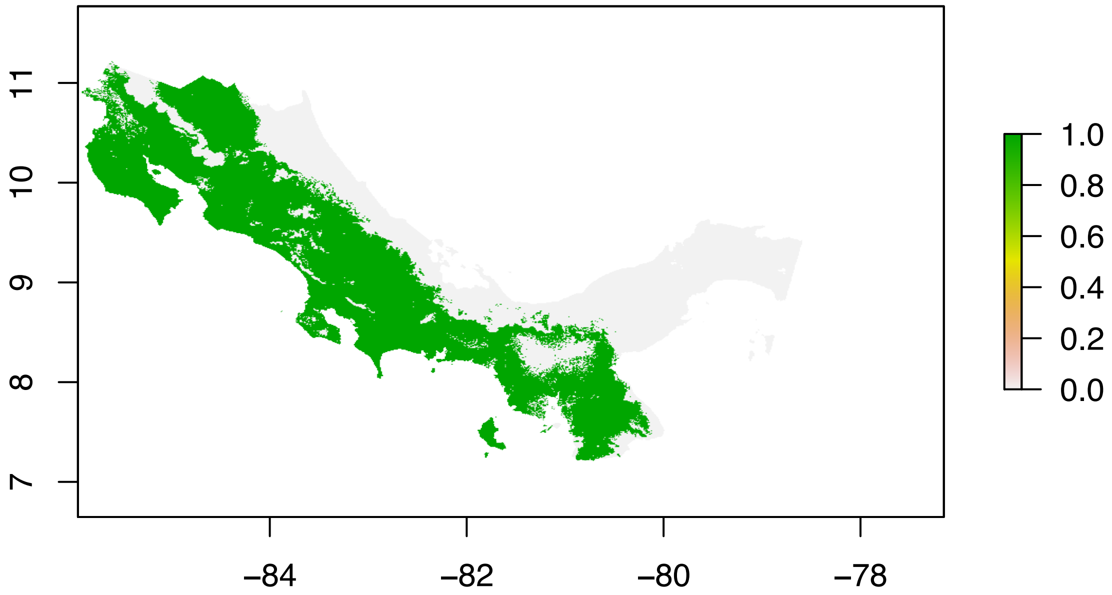


```
#100% threshold
plot(r_pred_100)
```

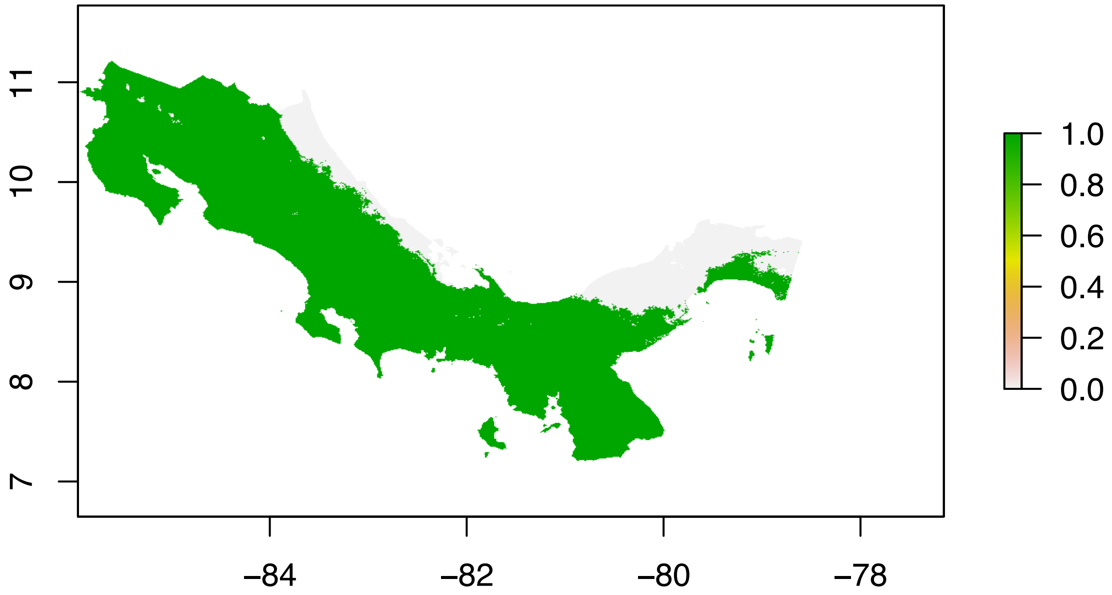


#### Section 5: Statistical analysis

**5.1 Make a presence-absence matrix for each threshold** The presence absence matrix tells us for the given locality (Madre Selva) which species the models predict to be there and which species it predicts to not be there. Since it is a binary prediction, 0 indicates an absence and 1 indicates a presence.
We have a presence absence matrix for each threshold because each one gives different predictions.

```
y=data.frame(LONGITUDE=-83.867218, LATITUDE=9.661011)

#90
maps=paste0(filepath,"90/",list.files(paste0(filepath,"output_trainingarea/90/"),pattern="*90.tif"))
x=stack(maps)
xy=raster::extract(x,y)
```

```
xy=t(xy)
#write.csv(xy,paste0(filepath, "output_trainingarea/PAM/PAM_90.csv"),quote=F)

#75
maps=paste0(filepath,"75/",list.files(paste0(filepath,"output_trainingarea/75/"),pattern="*75.tif"))
x=stack(maps)
xy=raster::extract(x,y)
xy=t(xy)
#write.csv(xy,paste0(filepath, "output_trainingarea/PAM/PAM_75.csv"),quote=F)

#100
maps=paste0(filepath,"100/",list.files(paste0(filepath,"output_trainingarea/100/"),pattern="*100.tif"))
x=stack(maps)
xy=raster::extract(x,y)
xy=t(xy)
#write.csv(xy,paste0(filepath, "output_trainingarea/PAM/PAM_100.csv"),quote=F)

#all points
maps=paste0(filepath,"all/",list.files(paste0(filepath,"output_trainingarea/all/"),pattern="*pts.tif"))
x=stack(maps)
xy=raster::extract(x,y)
xy=t(xy)
#write.csv(xy,paste0(filepath, "output_trainingarea/PAM/PAM_all.csv"),quote=F)
```

Here is an example of a presence absence matrix; I added the "recorded" column later

```
## # A tibble: 651 x 3
##    X1                         Predicted Recorded
##    <chr>                          <dbl>    <dbl>
##  1 Accipiter bicolor                  0        0
##  2 Accipiter cooperii                 1        0
##  3 Accipiter striatus                 0        0
##  4 Accipiter superciliosus            0        0
##  5 Actitis macularius                 0        0
##  6 Agelaius phoeniceus                0        0
##  7 Amaurolimnas concolor              0        0
##  8 Amazilia amabilis                  0        0
##  9 Amazilia boucardi                  0        0
## 10 Amazilia decora                    0        0
## # ... with 641 more rows
```

##### 5.2 Sorenson and Prediction Success Calculations (code from J.C. Cooper and J. Soberón)

By comparing the predicted presences and absences with the recorded presences and absences (our data), we can see how accurately the models predict the makeup of the ecosystem in Madre Selva.

```
alln=651

setwd(filepath)
files=paste0(filepath,"output_trainingarea/PAM/",list.files(paste0(filepath,"output_trainingarea/PAM/"))

for(i in 1:length(files)){
  x=files[i]
  csv=read.csv(x)
  n=nrow(csv)
```

```
  #Determine true presence from checklist
  actually.present=which(csv[,3]=="1")
  predicted.present=which(csv[,2]=="1")

  #Stats for no M
  #True Positives
  n.tp=length(which(actually.present%in%predicted.present))
  #True Negatives
  n.tn=alln-length(unique(c(actually.present,predicted.present)))
  #False Negative
  n.fn=length(actually.present)-n.tp
  #False Positive
  n.fp=length(predicted.present)-n.tp

  #Create a string to add to a matrix to save
  newrow=cbind(x,n.tp,n.fp,n.tn,n.fn)
  colnames(newrow)[1]="Locality"
  if(i==1){
    xarray=as.data.frame(newrow)
  }else{
    xarray=rbind(xarray,newrow)
  }
}

for(i in 2:ncol(xarray)){
  xarray[,i]=as.numeric(as.character(xarray[,i]))
}
```

**5.3 Calculate Sorensen and Chi-Square metrics** The prediction success is the correct predictions divided by all the predictions (tp + tn / all).
The Sorensen index is (2tp / 2tp + fp + fn); it is similar to the prediction success but weighs the presences more heavily than absences. This is useful because we do not have true absence data.

```
pred.success=function(tp,tn,sp){
  return((tp+tn)/sp)
}

sensitivity=function(tp,fn){
  return(tp/(tp+fn))
}

specificity=function(tn,fp){
  return(tn/(tn+fp))
}

Sorensen=function(tp,fn,fp){
  return((2*tp)/(2*tp+fn+fp))
}

for(i in 1:nrow(xarray)){
  #Calculate metrics for no M
  if(i==1){
    x2=pred.success(tp=xarray$n.tp[i],tn=xarray$n.tn[i],sp=alln)
```

```
    y2=Sorensen(tp=xarray$n.tp[i],fn=xarray$n.fn[i],fp=xarray$n.fp[i])
    z2=sensitivity(tp=xarray$n.tp[i],fn=xarray$n.fn[i])
    w2=specificity(tn=xarray$n.tn[i],fp=xarray$n.fp[i])

  }else{
    x=pred.success(tp=xarray$n.tp[i],tn=xarray$n.tn[i],sp=alln)
    y=Sorensen(tp=xarray$n.tp[i],fn=xarray$n.fn[i],fp=xarray$n.fp[i])
    z=sensitivity(tp=xarray$n.tp[i],fn=xarray$n.fn[i])
    w=specificity(tn=xarray$n.tn[i],fp=xarray$n.fp[i])
    x2=c(x2,x)
    y2=c(y2,y)
    z2=c(z2,z)
    w2=c(w2,w)
  }
}

Prediction.Success.NO=x2
Sorensen.NO=y2
sensitivity.NO=z2
specificity.NO=w2

xarray2=as.data.frame(cbind(xarray,Prediction.Success.NO,Sorensen.NO,sensitivity.NO,specificity.NO))

#write.csv(xarray2,paste0(paste0(filepath, "output_trainingarea/PAM/PAM.array2.csv"),quote=F)
```

```
## # A tibble: 3 x 10
##   Threshold  n.tp  n.fp  n.tn  n.fn total `Prediction suc~ `Sorensen index`
##       <dbl> <dbl> <dbl> <dbl> <dbl> <dbl>            <dbl>            <dbl>
## 1       100    76   286   289     0   651            0.561            0.347
## 2        75    56    23   552    20   651            0.934            0.723
## 3        90    65    50   525    11   651            0.906            0.681
## # ... with 2 more variables: Sensitivity <dbl>, Specificity <dbl>
```

**5.4 Plot** By plotting we can see that there is a tradeoff between sensitivity and specificity, where the 100% threshold model is better for sensitivity and the 75% threshold is better for specificity, but overall, thresholding at 75% is best for the models.

Generally, the prediction success is very high, which means that the observations acquired from eBird are a good source of data to make models. These are useful tests to perform because it is the underlying data that is used in a lot of ornithological research, and it's important to make sure that it is not too erroneous or biased.

```
PAM.array=read.csv(paste0(filepath, "output_real/PAM/PAM.array2.csv"))
subset=PAM.array[,c(1,6,7,8,9)]

PAM.array = read.csv(paste0(filepath,"output_real/PAM/PAM.array2.csv")) %>%
  select(Threshold, Sensitivity,Specificity,Prediction.success,Sorensen.index)%>%
  pivot_longer(cols=c(Sensitivity,Specificity,Prediction.success,Sorensen.index))%>%
  mutate(Threshold=as.character(Threshold)) %>%
  mutate(name = fct_relevel(name, "Sensitivity", "Specificity", "Prediction success", "Sorensen index")
  PAM.array$Threshold = factor(PAM.array$Threshold, levels = c("100", "90", "75"))
ggplot(PAM.array) +
  geom_col(aes(x=name,y=value,fill=Threshold),position = "dodge",) +
  xlab("Statistics") +
```

```r
  scale_fill_manual(values = c("#741B47", "#C27BA0", "#EAD1DC")) +
  labs(title = "Statistics for each threshold value")
```

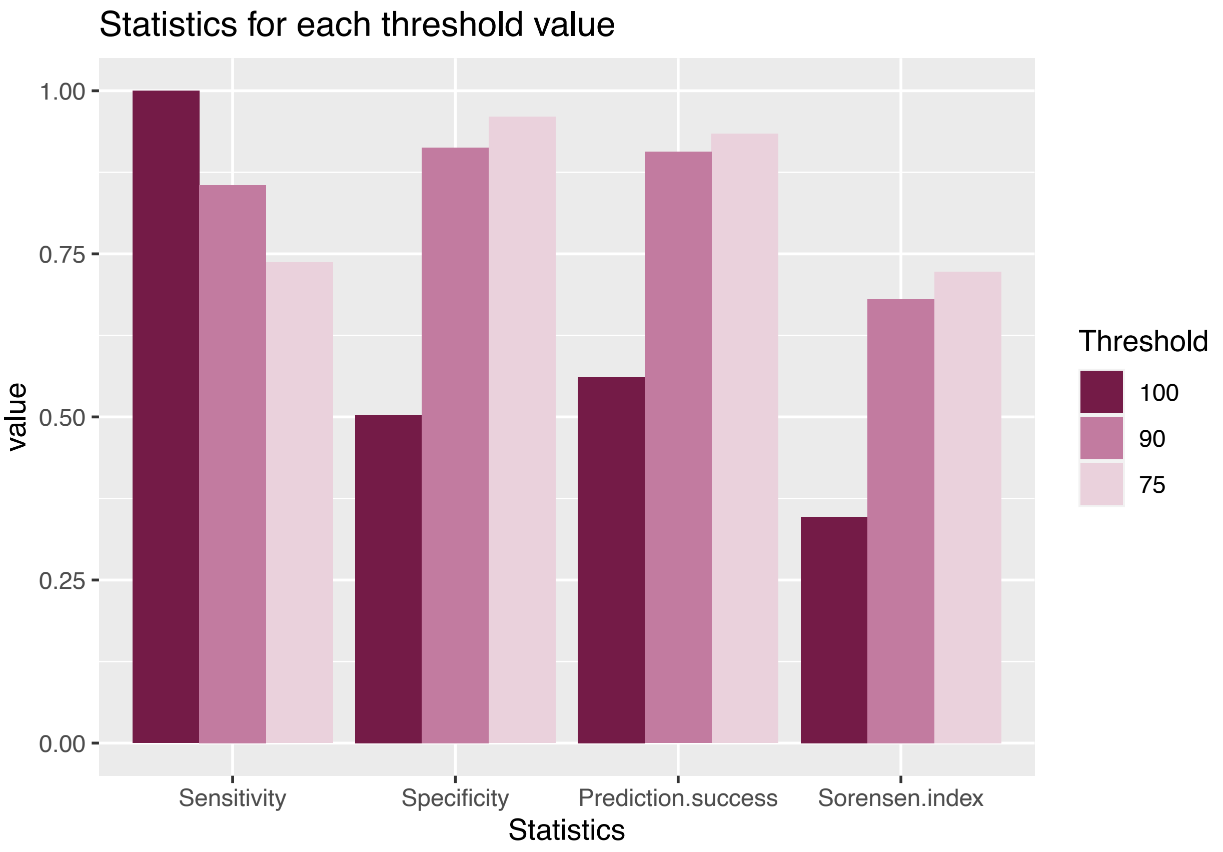


#### Section 6: Additional project for class: binning data

Since I had done all of this research over the summer, I decided to do an additional side project for the purpose of this class. Basically, I wanted to see if there was an ideal number of observations to make the models. If there are too few observations, we could predict a decrease in the quality of the models because there is not enough data to take into account all of the variables.
However for birds with too many observations, the quality could also decrease; a very common species is more likely to have vagrants that end up in areas that do not match their usual habitat, which could skew the models like we saw with the 100% thresholds.
I binned the species into categories based on the number of observations they had.

```r
library(lubridate)
library(gridExtra)
library(tidyverse)
library(data.table)
library(fossil)
library(geosphere)
library(rgdal)
library("brew")
library(ggplot2)
library(dplyr)
```

```
x_total = data.frame()
for (i in 1:length(species.files)){
  genus=strsplit(species.files[i],"-")[[1]][1]
  epithet=strsplit(species.files[i],"-")[[1]][2]
  species=strsplit(epithet,"[_]")[[1]][1]
  name = paste0(genus," ",species)
  filename=paste0(genus,"-",species,"_tobin.csv")
  file=read_csv(paste0(path,"artificial.absence/",species.files[i]))
  x= data.frame("species" = name, "observations" = paste0(nrow(file)))
  x_total <- rbind(x_total,x)
  #write.csv(x,paste0(SaveDir,filename),row.names = F)
}

#write.csv(x_total,paste0(SaveDir,"Species-observations_rownames.csv"))
```

##### 6.1 Make csv with species and # of observations used (from rarified)

**6.2 Bin species** I divided the data into about 6 even bins

```
observation.number=read_csv(paste0(SaveDir, "Species-observations.csv"))
observation.number=observation.number[order(observation.number$observations),]

Bin1.34=observation.number[1:110,]
#write.csv(Bin1.34,paste0(SaveDir, "/bins/Bin1.34.csv"))
Bin2.59=observation.number[111:222,]
#write.csv(Bin2.59,paste0(SaveDir, "/bins/Bin2.59.csv"))
Bin3.91=observation.number[223:326,]
#write.csv(Bin3.91,paste0(SaveDir, "/bins/Bin3.91.csv"))
Bin4.147=observation.number[327:434,]
#write.csv(Bin4.147,paste0(SaveDir, "/bins/Bin4.147.csv"))
Bin5.277=observation.number[435:548,]
#write.csv(Bin5.277,paste0(SaveDir, "/bins/Bin5.277.csv"))
Bin6.1378=observation.number[549:654,]
#write.csv(Bin6.1378,paste0(SaveDir, "/bins/Bin6.1378.csv"))
```

```
PAMall=read_csv(paste0(filepath, "output_real/PAM/PAM_75.csv"))
bins=paste0(filepath, "binning/bins/",list.files(paste0(filepath, "binning/decjan/"),pattern="*.csv"))
bins=unlist(bins)
for (i in 1:length(bins)){
  bin=read_csv(bins[i])
  names=as.vector(bin$species)
  Pam_Bin = PAMall %>% filter(X1 %in% names)
  name1=strsplit(bins[i],"/")[[1]][9]
  #write_csv(Pam_Bin,paste0(SaveDir, "/PAMs/",name1))
}
```

##### 6.3 Make PAMs

```
files=paste0(filepath,"/binning/decjan/", list.files(paste0(filepath, "/binning/decjan/"),pattern="*.cs

for(i in 1:length(files)){
```

```
  x=files[i]
  csv=read.csv(x)
  alln=nrow(csv)

  #Determine true presence from checklist
  actually.present=which(csv[,3]=="1")
  predicted.present=which(csv[,2]=="1")

  #Stats for no M
  #True Positives
  n.tp=length(which(actually.present%in%predicted.present))
  #True Negatives
  n.tn=alln-length(unique(c(actually.present,predicted.present)))
  #False Negative
  n.fn=length(actually.present)-n.tp
  #False Positive
  n.fp=length(predicted.present)-n.tp

  #Create a string to add to a matrix to save
  newrow=cbind(x,n.tp,n.fp,n.tn,n.fn)
  colnames(newrow)[1]="Locality"
  if(i==1){
    xarray=as.data.frame(newrow)
  }else{
    xarray=rbind(xarray,newrow)
  }
}

for(i in 2:ncol(xarray)){
  xarray[,i]=as.numeric(as.character(xarray[,i]))
}
```

##### 6.4 Get array and Sorensen index

```
pred.success=function(tp,tn,sp){
  return((tp+tn)/sp)
}

Sorensen=function(tp,fn,fp){
  return((2*tp)/(2*tp+fn+fp))
}

for(i in 1:nrow(xarray)){
  #Calculate metrics for no M
  if(i==1){
    x2=pred.success(tp=xarray$n.tp[i],tn=xarray$n.tn[i],sp=alln)
    y2=Sorensen(tp=xarray$n.tp[i],fn=xarray$n.fn[i],fp=xarray$n.fp[i])

  }else{
    x=pred.success(tp=xarray$n.tp[i],tn=xarray$n.tn[i],sp=alln)
    y=Sorensen(tp=xarray$n.tp[i],fn=xarray$n.fn[i],fp=xarray$n.fp[i])
    x2=c(x2,x)
    y2=c(y2,y)
```

```
  }
}

Prediction.Success.NO=x2
Sorensen.NO=y2

xarray2=as.data.frame(cbind(xarray,Prediction.Success.NO,Sorensen.NO))

#write.csv(xarray2,paste0(SaveDir, "PAMs/PAM.array.csv"),quote=F)

read_csv(paste0(filepath, "binning/decjan/PAM.array.csv"))
```

##### 6.5 Calculate Sorensen and Chi-Square metrics

```
## # A tibble: 6 x 8
##   Observations  n.tp  n.fp  n.tn  n.fn     n Prediction.success Sorensen.index
##   <chr>        <dbl> <dbl> <dbl> <dbl> <dbl>              <dbl>          <dbl>
## 1 10-34            8    13    85     1   107              0.869          0.533
## 2 35-59           20    13    74     5   112              0.839          0.690
## 3 60-91           16     9    77     2   104              0.894          0.744
## 4 92-147          17     3    86     2   108              0.954          0.872
## 5 148-277          8     4    99     3   114              0.939          0.696
## 6 278-1378        11     2    76    17   106              0.821          0.537
```

**6.6 Plot indices** We can see from the graph that the models had the best predictive success when there is a medium number of observations, in this case around 100-150 observations. The success rate of this bin is 1, which is extremely high. This supports my hypothesis that too many and too few data points can be detrimental.

```
PAM.bins=read.csv(paste0(filepath, "binning/decjan/PAM.array.csv"))
barplot(height=PAM.bins$Sorensen.index, names=PAM.bins$Observations,xlab="Number of Observations", ylab
```

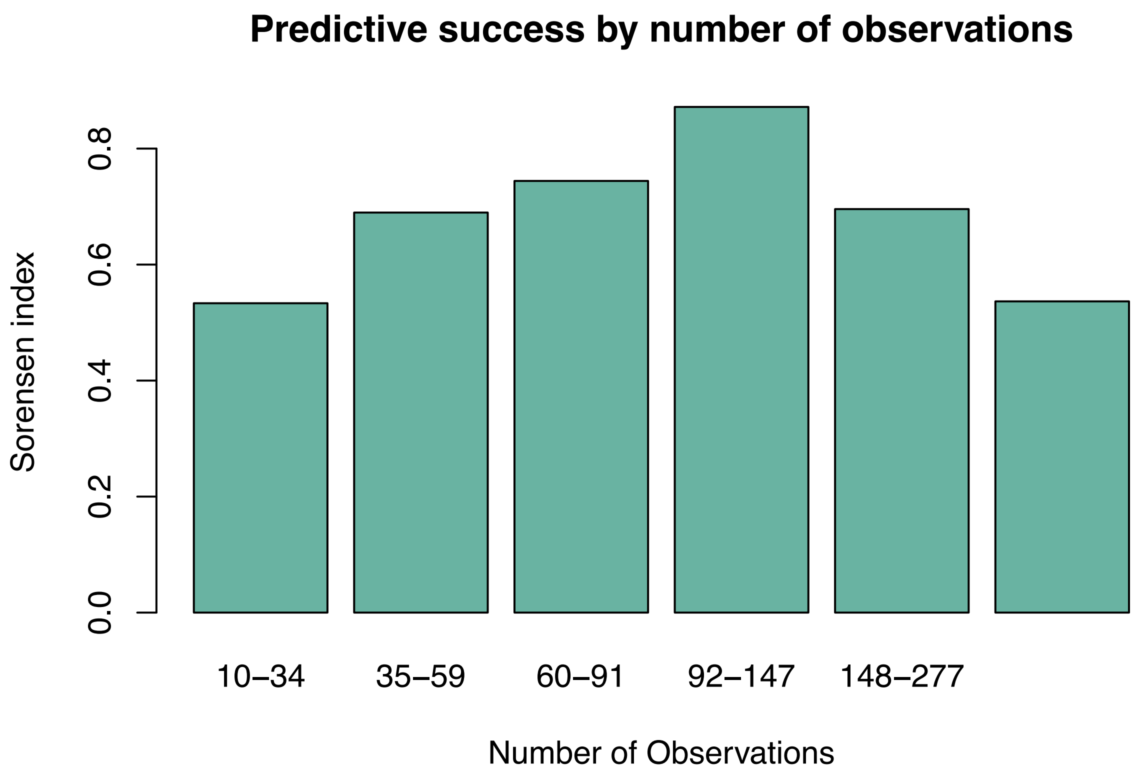
Predictive success by number of observations
Sorensen index
0.0
0.2
0.4
0.6
0.8
10–34
35–59
60–91
92–147
148–277
Number of Observations
