## Appendix 2 for "Testing the accuracy of species distribution models based on community science data"

Species and number of observations, winter Madre Selva

| species | observations |
|---|---|
| Accipiter bicolor | 30 |
| Accipiter cooperii | 24 |
| Accipiter striatus | 21 |
| Accipiter superciliosus | 25 |
| Actitis macularius | 526 |
| Agelaius phoeniceus | 109 |
| Amaurolimnas concolor | 13 |
| Amazilia amabilis | 80 |
| Amazilia boucardi | 38 |
| Amazilia decora | 92 |
| Amazilia edward | 127 |
| Amazilia hoffmanni | 193 |
| Amazilia rutila | 146 |
| Amazilia tzacatl | 856 |
| Amazona albifrons | 243 |
| Amazona auropalliata | 88 |
| Amazona autumnalis | 441 |
| Amazona farinosa | 226 |
| Amblycercus holosericeus | 93 |
| Anabacerthia variegaticeps | 20 |
| Anhinga anhinga | 200 |
| Anthracothorax prevostii | 190 |
| Antrostomus carolinensis | 19 |
| Ara macao | 223 |
| Aramides albiventris | 81 |
| Aramides cajaneus | 169 |
| Aramus guarauna | 42 |
| Archilochus colubris | 208 |

Species and number of observations, summer San José

| species | observations |
|---|---|
| Acanthidops bairdi | 16 |
| Accipiter bicolor | 13 |
| Actitis macularius | 12 |
| Agelaius phoeniceus | 105 |
| Amaurolimnas concolor | 12 |
| Amazilia amabilis | 70 |
| Amazilia hoffmanni | 144 |
| Amazilia rutila | 102 |
| Amazilia tzacatl | 622 |
| Amazona albifrons | 135 |
| Amazona auropalliata | 59 |
| Amazona autumnalis | 315 |
| Amazona farinosa | 161 |
| Amblycercus holosericeus | 76 |
| Anabacerthia variegaticeps | 12 |
| Anhinga anhinga | 146 |
| Anthracothorax prevostii | 72 |
| Aphanotriccus capitalis | 10 |
| Apodidae sp | 53 |
| Ara macao | 140 |
| Aramides albiventris | 68 |
| Aramides cajaneus | 161 |
| Aramus guarauna | 31 |
| Ardea alba | 307 |
| Ardea herodias | 48 |
| Arremon aurantiirostris | 181 |
| Arremon brunneinucha | 89 |
| Arremon crassirostris | 20 |

| | |
|---|---|
| **Ardea alba** | 586 |
| **Ardea herodias** | 326 |
| **Arenaria interpres** | 75 |
| **Arremon aurantiirostris** | 239 |
| **Arremon brunneinucha** | 98 |
| **Arremon costaricensis** | 14 |
| **Arremon crassirostris** | 26 |
| **Arremonops conirostris** | 322 |
| **Arremonops rufivirgatus** | 49 |
| **Asio clamator** | 14 |
| **Atalotriccus pilaris** | 21 |
| **Atlapetes albinucha** | 89 |
| **Atlapetes tibialis** | 70 |
| **Attila spadiceus** | 294 |
| **Aulacorhynchus prasinus** | 127 |
| **Automolus exsertus** | 47 |
| **Automolus ochrolaemus** | 49 |
| **Automolus subulatus** | 20 |
| **Aythya affinis** | 55 |
| **Aythya collaris** | 13 |
| **Bangsia arcaei** | 30 |
| **Baryphthengus martii** | 105 |
| **Basileuterus culicivorus** | 105 |
| **Basileuterus melanogenys** | 52 |
| **Basileuterus melanotis** | 46 |
| **Basileuterus rufifrons** | 308 |
| **Bolborhynchus lineola** | 46 |
| **Brotogeris jugularis** | 725 |
| **Bubulcus ibis** | 731 |
| **Burhinus bistriatus** | 64 |
| **Buteo albonotatus** | 126 |
| **Buteo brachyurus** | 326 |
| **Buteo jamaicensis** | 104 |

| | |
|---|---|
| **Arremonops conirostris** | 302 |
| **Arremonops rufivirgatus** | 84 |
| **Asio clamator** | 13 |
| **Atlapetes albinucha** | 71 |
| **Atlapetes tibialis** | 63 |
| **Attila spadiceus** | 228 |
| **Aulacorhynchus prasinus** | 98 |
| **Automolus ochrolaemus** | 43 |
| **Automolus subulatus** | 11 |
| **Bangsia arcaei** | 14 |
| **Baryphthengus martii** | 109 |
| **Basileuterus culicivorus** | 102 |
| **Basileuterus melanogenys** | 54 |
| **Basileuterus melanotis** | 43 |
| **Basileuterus rufifrons** | 253 |
| **Bolborhynchus lineola** | 19 |
| **Brotogeris jugularis** | 619 |
| **Bubulcus ibis** | 386 |
| **Buteo albonotatus** | 47 |
| **Buteo brachyurus** | 144 |
| **Buteo jamaicensis** | 52 |
| **Buteo plagiatus** | 178 |
| **Buteogallus anthracinus** | 166 |
| **Buteogallus urubitinga** | 44 |
| **Butorides striata** | 34 |
| **Butorides virescens** | 278 |
| **Cacicus uropygialis** | 110 |
| **Cairina moschata** | 97 |
| **Calocitta formosa** | 155 |
| **Campephilus guatemalensis** | 137 |
| **Camptostoma imberbe** | 31 |
| **Camptostoma obsoletum** | 63 |
| **Campylopterus hemileucuru** | 107 |

| | |
|---|---|
| **Buteo nitidus** | 85 |
| **Buteo plagiatus** | 326 |
| **Buteo platypterus** | 504 |
| **Buteo swainsoni** | 59 |
| **Buteogallus anthracinus** | 278 |
| **Buteogallus urubitinga** | 60 |
| **Butorides virescens** | 384 |
| **Cacicus uropygialis** | 159 |
| **Cairina moschata** | 88 |
| **Calidris alba** | 72 |
| **Calidris himantopus** | 12 |
| **Calidris mauri** | 35 |
| **Calidris minutilla** | 92 |
| **Calidris pusilla** | 41 |
| **Calocitta formosa** | 198 |
| **Campephilus guatemalensis** | 243 |
| **Camptostoma imberbe** | 38 |
| **Camptostoma obsoletum** | 147 |
| **Campylopterus hemileucuru** | 133 |
| **Campylorhamphus pusillus** | 49 |
| **Campylorhynchus rufinucha** | 310 |
| **Campylorhynchus zonatus** | 63 |
| **Cantorchilus elutus** | 161 |
| **Cantorchilus modestus** | 222 |
| **Cantorchilus nigricapillus** | 243 |
| **Cantorchilus semibadius** | 156 |
| **Cantorchilus thoracicus** | 112 |
| **Cantorchilus zeledoni** | 37 |
| **Capsiempis flaveola** | 175 |
| **Caracara cheriway** | 519 |
| **Cardellina canadensis** | 32 |
| **Cardellina pusilla** | 319 |
| **Carpodectes antoniae** | 16 |

| | |
|---|---|
| **Campylorhamphus pusillus** | 29 |
| **Campylorhynchus rufinucha** | 271 |
| **Campylorhynchus zonatus** | 42 |
| **Cantorchilus modestus** | 208 |
| **Cantorchilus nigricapillus** | 148 |
| **Cantorchilus thoracicus** | 103 |
| **Cantorchilus zeledoni** | 20 |
| **Capsiempis flaveola** | 110 |
| **Caracara cheriway** | 256 |
| **Carpodectes nitidus** | 12 |
| **Caryothraustes poliogaster** | 62 |
| **Cathartes aura** | 893 |
| **Cathartes burrovianus** | 40 |
| **Catharus aurantiirostris** | 173 |
| **Catharus frantzii** | 68 |
| **Catharus fuscater** | 79 |
| **Catharus gracilirostris** | 52 |
| **Catharus mexicanus** | 39 |
| **Celeus castaneus** | 11 |
| **Celeus loricatus** | 72 |
| **Ceratopipra mentalis** | 86 |
| **Cercomacroides tyrannina** | 100 |
| **Chaetura cinereiventris** | 42 |
| **Chaetura fumosa** | 37 |
| **Chaetura sp** | 19 |
| **Chaetura vauxi** | 136 |
| **Chalybura urochrysia** | 59 |
| **Chamaepetes unicolor** | 63 |
| **Charadrius vociferus** | 10 |
| **Chiroxiphia linearis** | 151 |
| **Chloroceryle aenea** | 53 |
| **Chloroceryle amazona** | 174 |
| **Chloroceryle americana** | 238 |

| | |
|---|---|
| **Carpodectes nitidus** | 25 |
| **Caryothraustes poliogaster** | 58 |
| **Cathartes aura** | 1329 |
| **Cathartes burrovianus** | 43 |
| **Catharus aurantiirostris** | 108 |
| **Catharus frantzii** | 52 |
| **Catharus fuscater** | 65 |
| **Catharus gracilirostris** | 36 |
| **Catharus mexicanus** | 42 |
| **Catharus minimus** | 12 |
| **Catharus ustulatus** | 172 |
| **Celeus castaneus** | 25 |
| **Celeus loricatus** | 75 |
| **Cephalopterus glabricollis** | 20 |
| **Ceratopipra mentalis** | 148 |
| **Cercomacroides tyrannina** | 163 |
| **Chaetura cinereiventris** | 55 |
| **Chaetura fumosa** | 90 |
| **Chaetura vauxi** | 174 |
| **Chalybura urochrysia** | 89 |
| **Chamaepetes unicolor** | 76 |
| **Charadrius collaris** | 28 |
| **Charadrius semipalmatus** | 86 |
| **Charadrius vociferus** | 51 |
| **Charadrius wilsonia** | 48 |
| **Chiroxiphia lanceolata** | 58 |
| **Chiroxiphia linearis** | 145 |
| **Chlidonias niger** | 59 |
| **Chloroceryle aenea** | 74 |
| **Chloroceryle amazona** | 242 |
| **Chloroceryle americana** | 316 |
| **Chloroceryle inda** | 12 |
| **Chlorophanes spiza** | 295 |

| | |
|---|---|
| **Chloroceryle inda** | 11 |
| **Chlorophanes spiza** | 203 |
| **Chlorophonia callophrys** | 101 |
| **Chlorospingus flavopectus** | 170 |
| **Chlorospingus pileatus** | 66 |
| **Chlorostilbon assimilis** | 73 |
| **Chlorostilbon canivetii** | 92 |
| **Chlorothraupis carmioli** | 57 |
| **Chondrohierax uncinatus** | 27 |
| **Chordeiles acutipennis** | 24 |
| **Chordeiles minor** | 23 |
| **Chrysothlypis chrysomelas** | 47 |
| **Ciccaba nigrolineata** | 27 |
| **Ciccaba virgata** | 44 |
| **Cinclus mexicanus** | 27 |
| **Claravis pretiosa** | 98 |
| **Cochlearius cochlearius** | 88 |
| **Coereba flaveola** | 362 |
| **Colaptes rubiginosus** | 103 |
| **Colibri cyanotus** | 75 |
| **Colibri delphinae** | 37 |
| **Colinus cristatus** | 71 |
| **Colonia colonus** | 86 |
| **Columba livia** | 114 |
| **Columbina inca** | 286 |
| **Columbina minuta** | 70 |
| **Columbina passerina** | 149 |
| **Columbina talpacoti** | 576 |
| **Conopias albovittatus** | 43 |
| **Contopus cinereus** | 157 |
| **Contopus lugubris** | 40 |
| **Coragyps atratus** | 1041 |
| **Corapipo altera** | 69 |

| | |
|---|---|
| **Chlorophonia callophrys** | 117 |
| **Chlorospingus flavopectus** | 197 |
| **Chlorospingus pileatus** | 67 |
| **Chlorostilbon assimilis** | 98 |
| **Chlorostilbon canivetii** | 103 |
| **Chlorothraupis carmioli** | 78 |
| **Chondrohierax uncinatus** | 64 |
| **Chordeiles acutipennis** | 78 |
| **Chrysothlypis chrysomelas** | 66 |
| **Ciccaba nigrolineata** | 41 |
| **Ciccaba virgata** | 85 |
| **Cinclus mexicanus** | 31 |
| **Claravis pretiosa** | 117 |
| **Coccyzus americanus** | 30 |
| **Cochlearius cochlearius** | 119 |
| **Coereba flaveola** | 542 |
| **Colaptes rubiginosus** | 141 |
| **Colibri cyanotus** | 115 |
| **Colibri delphinae** | 42 |
| **Colinus cristatus** | 30 |
| **Colonia colonus** | 111 |
| **Columba livia** | 194 |
| **Columbina inca** | 406 |
| **Columbina minuta** | 79 |
| **Columbina passerina** | 183 |
| **Columbina talpacoti** | 697 |
| **Conopias albovittatus** | 51 |
| **Contopus cinereus** | 241 |
| **Contopus cooperi** | 58 |
| **Contopus lugubris** | 42 |
| **Contopus sordidulus** | 64 |
| **Contopus virens** | 98 |
| **Coragyps atratus** | 1378 |

| | |
|---|---|
| **Cranioleuca erythrops** | 68 |
| **Crax rubra** | 84 |
| **Crotophaga ani** | 215 |
| **Crotophaga sulcirostris** | 406 |
| **Crypturellus soui** | 198 |
| **Cyanerpes cyaneus** | 341 |
| **Cyanerpes lucidus** | 119 |
| **Cyanoloxia cyanoides** | 156 |
| **Cyanolyca cucullata** | 14 |
| **Cyclarhis gujanensis** | 99 |
| **Cymbilaimus lineatus** | 46 |
| **Cyphorhinus phaeocephalus** | 46 |
| **Cypseloides cherriei** | 10 |
| **Cypseloides niger** | 25 |
| **Dacnis cayana** | 109 |
| **Dacnis venusta** | 138 |
| **Dendrocincla fuliginosa** | 57 |
| **Dendrocincla homochroa** | 45 |
| **Dendrocolaptes sanctithoma** | 111 |
| **Dendrocolaptinae sp** | 32 |
| **Dendrocygna autumnalis** | 303 |
| **Dendrortyx leucophrys** | 19 |
| **Diglossa plumbea** | 76 |
| **Discosura conversii** | 33 |
| **Dives dives** | 383 |
| **Doryfera ludovicae** | 22 |
| **Dryobates fumigatus** | 70 |
| **Dryobates villosus** | 52 |
| **Dryocopus lineatus** | 255 |
| **Dysithamnus mentalis** | 50 |
| **Dysithamnus striaticeps** | 16 |
| **Egretta caerulea** | 187 |
| **Egretta thula** | 151 |

| | |
|---|---|
| **Corapipo altera** | 182 |
| **Cotinga ridgwayi** | 27 |
| **Cranioleuca erythrops** | 101 |
| **Crax rubra** | 108 |
| **Crotophaga ani** | 222 |
| **Crotophaga sulcirostris** | 525 |
| **Crypturellus soui** | 191 |
| **Cyanerpes cyaneus** | 396 |
| **Cyanerpes lucidus** | 165 |
| **Cyanocorax affinis** | 90 |
| **Cyanoloxia cyanoides** | 191 |
| **Cyanolyca cucullata** | 28 |
| **Cyclarhis gujanensis** | 96 |
| **Cymbilaimus lineatus** | 79 |
| **Cyphorhinus phaeocephalus** | 77 |
| **Cypseloides cherriei** | 19 |
| **Cypseloides niger** | 14 |
| **Dacnis cayana** | 213 |
| **Dacnis venusta** | 140 |
| **Deconychura longicauda** | 43 |
| **Dendrocincla anabatina** | 39 |
| **Dendrocincla fuliginosa** | 77 |
| **Dendrocincla homochroa** | 48 |
| **Dendrocolaptes sanctithoma** | 149 |
| **Dendrocygna autumnalis** | 184 |
| **Dendrortyx leucophrys** | 19 |
| **Diglossa plumbea** | 87 |
| **Discosura conversii** | 65 |
| **Dives dives** | 426 |
| **Dixiphia pipra** | 22 |
| **Doryfera ludovicae** | 20 |
| **Dryobates fumigatus** | 107 |
| **Dryobates villosus** | 68 |

| | |
|---|---|
| **Egretta tricolor** | 75 |
| **Elaenia chiriquensis** | 83 |
| **Elaenia flavogaster** | 438 |
| **Elaenia frantzii** | 125 |
| **Elanoides forficatus** | 248 |
| **Elanus leucurus** | 98 |
| **Electron carinatum** | 18 |
| **Electron platyrhynchum** | 107 |
| **Elvira cupreiceps** | 33 |
| **Empidonax atriceps** | 31 |
| **Empidonax flavescens** | 82 |
| **Epinecrophylla fulviventris** | 39 |
| **Eubucco bourcierii** | 38 |
| **Eucometis penicillata** | 113 |
| **Eudocimus albus** | 209 |
| **Eugenes spectabilis** | 49 |
| **Eumomota superciliosa** | 179 |
| **Eupherusa eximia** | 58 |
| **Eupherusa nigriventris** | 22 |
| **Euphonia affinis** | 94 |
| **Euphonia anneae** | 71 |
| **Euphonia elegantissima** | 51 |
| **Euphonia gouldi** | 111 |
| **Euphonia hirundinacea** | 190 |
| **Euphonia luteicapilla** | 232 |
| **Euphonia minuta** | 41 |
| **Eupsittula canicularis** | 143 |
| **Eupsittula nana** | 69 |
| **Eurypyga helias** | 49 |
| **Eutoxeres aquila** | 22 |
| **Falco rufigularis** | 139 |
| **Florisuga mellivora** | 148 |
| **Formicarius analis** | 112 |

| | |
|---|---|
| **Dryocopus lineatus** | 432 |
| **Dumetella carolinensis** | 41 |
| **Dysithamnus mentalis** | 85 |
| **Dysithamnus puncticeps** | 25 |
| **Dysithamnus striaticeps** | 30 |
| **Egretta caerulea** | 473 |
| **Egretta thula** | 417 |
| **Egretta tricolor** | 181 |
| **Elaenia chiriquensis** | 62 |
| **Elaenia flavogaster** | 450 |
| **Elaenia frantzii** | 128 |
| **Elanoides forficatus** | 39 |
| **Elanus leucurus** | 206 |
| **Electron platyrhynchum** | 122 |
| **Elvira chionura** | 40 |
| **Elvira cupreiceps** | 56 |
| **Empidonax atriceps** | 27 |
| **Empidonax flavescens** | 108 |
| **Empidonax flaviventris** | 261 |
| **Empidonax minimus** | 31 |
| **Empidonax traillii** | 23 |
| **Empidonax virescens** | 97 |
| **Epinecrophylla fulviventris** | 71 |
| **Eubucco bourcierii** | 65 |
| **Eucometis penicillata** | 116 |
| **Eudocimus albus** | 236 |
| **Eugenes spectabilis** | 57 |
| **Eumomota superciliosa** | 132 |
| **Eupherusa eximia** | 77 |
| **Eupherusa nigriventris** | 22 |
| **Euphonia affinis** | 115 |
| **Euphonia anneae** | 117 |
| **Euphonia elegantissima** | 57 |

| | |
|---|---|
| **Fregata magnificens** | 267 |
| **Galbula ruficauda** | 78 |
| **Gallinula galeata** | 27 |
| **Geothlypis poliocephala** | 160 |
| **Geothlypis semiflava** | 59 |
| **Geotrygon montana** | 53 |
| **Glaucidium brasilianum** | 62 |
| **Glaucidium costaricanum** | 11 |
| **Glaucidium griseiceps** | 15 |
| **Glaucis aeneus** | 39 |
| **Glyphorynchus spirurus** | 139 |
| **Gymnocichla nudiceps** | 10 |
| **Gymnopithys bicolor** | 70 |
| **Habia fuscicauda** | 83 |
| **Habia rubica** | 75 |
| **Hafferia zeledoni** | 25 |
| **Harpagus bidentatus** | 72 |
| **Heliodoxa jacula** | 72 |
| **Heliomaster constantii** | 43 |
| **Heliomaster longirostris** | 81 |
| **Heliothryx barroti** | 125 |
| **Henicorhina leucophrys** | 166 |
| **Henicorhina leucosticta** | 200 |
| **Herpetotheres cachinnans** | 225 |
| **Himantopus mexicanus** | 49 |
| **Hylocharis eliciae** | 72 |
| **Hylopezus dives** | 39 |
| **Hylopezus perspicillatus** | 24 |
| **Hylophylax naevioides** | 65 |
| **Ibycter americanus** | 19 |
| **Icterus mesomelas** | 17 |
| **Icterus pectoralis** | 24 |
| **Icterus prosthemelas** | 97 |

| | |
|---|---|
| **Euphonia gouldi** | 156 |
| **Euphonia hirundinacea** | 280 |
| **Euphonia imitans** | 108 |
| **Euphonia laniirostris** | 167 |
| **Euphonia luteicapilla** | 429 |
| **Euphonia minuta** | 91 |
| **Eupsittula canicularis** | 215 |
| **Eupsittula nana** | 110 |
| **Eupsittula pertinax** | 56 |
| **Eurypyga helias** | 57 |
| **Eutoxeres aquila** | 36 |
| **Falco columbarius** | 46 |
| **Falco peregrinus** | 113 |
| **Falco rufigularis** | 163 |
| **Falco sparverius** | 108 |
| **Florisuga mellivora** | 226 |
| **Formicarius analis** | 108 |
| **Fregata magnificens** | 421 |
| **Fulica americana** | 35 |
| **Galbula ruficauda** | 131 |
| **Gallinago delicata** | 12 |
| **Gallinula galeata** | 42 |
| **Gampsonyx swainsonii** | 43 |
| **Gelochelidon nilotica** | 29 |
| **Geothlypis formosa** | 96 |
| **Geothlypis philadelphia** | 194 |
| **Geothlypis poliocephala** | 165 |
| **Geothlypis semiflava** | 70 |
| **Geothlypis tolmiei** | 16 |
| **Geothlypis trichas** | 32 |
| **Geotrygon montana** | 62 |
| **Geranospiza caerulescens** | 67 |
| **Glaucidium brasilianum** | 115 |

| | |
|---|---|
| **Icterus pustulatus** | 103 |
| **Ictinia plumbea** | 65 |
| **Ixobrychus exilis** | 10 |
| **Ixothraupis guttata** | 57 |
| **Jabiru mycteria** | 13 |
| **Jacana spinosa** | 156 |
| **Klais guimeti** | 85 |
| **Lampornis calolaemus** | 84 |
| **Lampornis castaneoventris** | 39 |
| **Lampornis hemileucus** | 13 |
| **Lanio leucothorax** | 26 |
| **Laterallus albigularis** | 105 |
| **Laterallus exilis** | 10 |
| **Legatus leucophaius** | 342 |
| **Leistes militaris** | 85 |
| **Lepidocolaptes affinis** | 49 |
| **Lepidocolaptes souleyetii** | 358 |
| **Leptodon cayanensis** | 44 |
| **Leptopogon amaurocephalu** | 18 |
| **Leptopogon superciliaris** | 47 |
| **Leptotila cassinii** | 89 |
| **Leptotila plumbeiceps** | 22 |
| **Leptotila verreauxi** | 633 |
| **Leptotrygon veraguensis** | 12 |
| **Leucophaeus atricilla** | 54 |
| **Leucopternis semiplumbeus** | 21 |
| **Lipaugus unirufus** | 49 |
| **Lophornis helenae** | 24 |
| **Lophostrix cristata** | 26 |
| **Lophotriccus pileatus** | 115 |
| **Malacoptila panamensis** | 55 |
| **Manacus candei** | 133 |
| **Margarornis rubiginosus** | 51 |

| | |
|---|---|
| **Glaucidium griseiceps** | 18 |
| **Glaucis aeneus** | 65 |
| **Glyphorynchus spirurus** | 233 |
| **Gymnocichla nudiceps** | 34 |
| **Gymnopithys bicolor** | 104 |
| **Habia fuscicauda** | 134 |
| **Habia rubica** | 87 |
| **Haematopus palliatus** | 37 |
| **Hafferia zeledoni** | 54 |
| **Harpagus bidentatus** | 160 |
| **Heliodoxa jacula** | 79 |
| **Heliomaster constantii** | 74 |
| **Heliomaster longirostris** | 102 |
| **Heliothryx barroti** | 212 |
| **Helmitheros vermivorum** | 54 |
| **Henicorhina leucophrys** | 196 |
| **Henicorhina leucosticta** | 237 |
| **Herpetotheres cachinnans** | 347 |
| **Heterospingus rubrifrons** | 14 |
| **Himantopus mexicanus** | 92 |
| **Hirundo rustica** | 382 |
| **Hylocharis eliciae** | 192 |
| **Hylocichla mustelina** | 151 |
| **Hylopezus dives** | 40 |
| **Hylopezus perspicillatus** | 23 |
| **Hylophilus flavipes** | 73 |
| **Hylophylax naevioides** | 91 |
| **Icterus galbula** | 797 |
| **Icterus pectoralis** | 22 |
| **Icterus prosthemelas** | 142 |
| **Icterus pustulatus** | 106 |
| **Icterus spurius** | 140 |
| **Ixothraupis guttata** | 86 |

| | |
|---|---|
| **Megaceryle torquata** | 265 |
| **Megarynchus pitangua** | 479 |
| **Megascops choliba** | 30 |
| **Megascops clarkii** | 12 |
| **Megascops cooperi** | 12 |
| **Melanerpes formicivorus** | 53 |
| **Melanerpes hoffmannii** | 403 |
| **Melanerpes pucherani** | 230 |
| **Melozone cabanisi** | 18 |
| **Melozone leucotis** | 72 |
| **Mesembrinibis cayennensis** | 54 |
| **Micrastur ruficollis** | 25 |
| **Micrastur semitorquatus** | 92 |
| **Microbates cinereiventris** | 30 |
| **Microcerculus marginatus** | 78 |
| **Microcerculus philomela** | 55 |
| **Microchera albocoronata** | 29 |
| **Microrhopias quixensis** | 88 |
| **Milvago chimachima** | 347 |
| **Mimus gilvus** | 172 |
| **Mionectes oleagineus** | 143 |
| **Mionectes olivaceus** | 57 |
| **Mitrephanes phaeocercus** | 69 |
| **Mitrospingus cassinii** | 35 |
| **Molothrus aeneus** | 269 |
| **Molothrus bonariensis** | 84 |
| **Molothrus oryzivorus** | 51 |
| **Momotus lessonii** | 285 |
| **Monasa morphoeus** | 19 |
| **Morococcyx erythropygus** | 66 |
| **Morphnarchus princeps** | 38 |
| **Myadestes melanops** | 108 |
| **Mycteria americana** | 134 |

| | |
|---|---|
| Jacana jacana | 72 |
| Jacana spinosa | 242 |
| Junco vulcani | 11 |
| Klais guimeti | 137 |
| Lampornis calolaemus | 99 |
| Lampornis castaneoventris | 48 |
| Lampornis hemileucus | 25 |
| Lanio leucothorax | 56 |
| Laterallus albigularis | 220 |
| Laterallus exilis | 21 |
| Leiothlypis peregrina | 782 |
| Leistes militaris | 77 |
| Lepidocolaptes affinis | 64 |
| Lepidocolaptes souleyetii | 441 |
| Lepidothrix coronata | 101 |
| Leptodon cayanensis | 90 |
| Leptopogon amaurocephalu | 18 |
| Leptopogon superciliaris | 72 |
| Leptotila cassinii | 113 |
| Leptotila plumbeiceps | 25 |
| Leptotila verreauxi | 710 |
| Leptotrygon veraguensis | 13 |
| Leucophaeus atricilla | 200 |
| Leucophaeus pipixcan | 38 |
| Leucopternis semiplumbeus | 28 |
| Limnodromus griseus | 31 |
| Lipaugus unirufus | 93 |
| Lonchura malacca | 15 |
| Lophornis adorabilis | 34 |
| Lophornis helenae | 44 |
| Lophostrix cristata | 44 |
| Lophotriccus pileatus | 156 |
| Lurocalis semitorquatus | 23 |

| | |
|---|---|
| Myiarchus nuttingi | 63 |
| Myiarchus tuberculifer | 316 |
| Myiarchus tyrannulus | 95 |
| Myiobius sulphureipygius | 57 |
| Myioborus miniatus | 149 |
| Myioborus torquatus | 68 |
| Myiodynastes hemichrysus | 42 |
| Myiodynastes luteiventris | 214 |
| Myiodynastes maculatus | 292 |
| Myiopagis viridicata | 70 |
| Myiornis atricapillus | 29 |
| Myiothlypis fulvicauda | 172 |
| Myiozetetes granadensis | 190 |
| Myiozetetes similis | 660 |
| Myrmotherula axillaris | 44 |
| Myrmotherula schisticolor | 58 |
| Notharchus hyperrhynchus | 72 |
| Notharchus tectus | 36 |
| Nothocercus bonapartei | 20 |
| Numenius phaeopus | 51 |
| Nyctanassa violacea | 68 |
| Nyctibius grandis | 33 |
| Nyctibius griseus | 35 |
| Nycticorax nycticorax | 61 |
| Nyctidromus albicollis | 151 |
| Odontophorus guttatus | 23 |
| Odontophorus leucolaemus | 23 |
| Oncostoma cinereigulare | 60 |
| Onychorhynchus coronatus | 42 |
| Oreothlypis gutturalis | 46 |
| Ornithion brunneicapillus | 36 |
| Ornithion semiflavum | 18 |
| Ortalis cinereiceps | 315 |

| | |
|---|---|
| **Malacoptila panamensis** | 79 |
| **Manacus aurantiacus** | 90 |
| **Manacus candei** | 166 |
| **Manacus vitellinus** | 83 |
| **Margarornis rubiginosus** | 58 |
| **Megaceryle alcyon** | 109 |
| **Megaceryle torquata** | 368 |
| **Megarynchus pitangua** | 628 |
| **Megascops choliba** | 35 |
| **Megascops clarkii** | 16 |
| **Megascops cooperi** | 39 |
| **Megascops guatemalae** | 13 |
| **Melanerpes chrysauchen** | 92 |
| **Melanerpes formicivorus** | 62 |
| **Melanerpes hoffmannii** | 508 |
| **Melanerpes pucherani** | 306 |
| **Melanerpes rubricapillus** | 408 |
| **Melozone cabanisi** | 22 |
| **Melozone leucotis** | 70 |
| **Mesembrinibis cayennensis** | 93 |
| **Micrastur ruficollis** | 56 |
| **Micrastur semitorquatus** | 104 |
| **Microbates cinereiventris** | 43 |
| **Microcerculus marginatus** | 99 |
| **Microcerculus philomela** | 54 |
| **Microchera albocoronata** | 38 |
| **Microrhopias quixensis** | 129 |
| **Milvago chimachima** | 524 |
| **Mimus gilvus** | 244 |
| **Mionectes oleagineus** | 227 |
| **Mionectes olivaceus** | 119 |
| **Mitrephanes phaeocercus** | 85 |
| **Mitrospingus cassinii** | 50 |

| | |
|---|---|
| **Pachyramphus aglaiae** | 124 |
| **Pachyramphus cinnamomeu** | 129 |
| **Pachyramphus polychopteru** | 136 |
| **Pachyramphus versicolor** | 28 |
| **Pachysylvia decurtata** | 392 |
| **Pandion haliaetus** | 80 |
| **Panterpe insignis** | 52 |
| **Panyptila cayennensis** | 33 |
| **Parabuteo unicinctus** | 14 |
| **Passer domesticus** | 101 |
| **Passerina caerulea** | 80 |
| **Patagioenas cayennensis** | 265 |
| **Patagioenas fasciata** | 113 |
| **Patagioenas flavirostris** | 366 |
| **Patagioenas nigrirostris** | 223 |
| **Patagioenas speciosa** | 98 |
| **Patagioenas subvinacea** | 58 |
| **Pelecanus occidentalis** | 240 |
| **Penelope purpurascens** | 171 |
| **Peucaea ruficauda** | 127 |
| **Pezopetes capitalis** | 31 |
| **Phaenostictus mcleannani** | 38 |
| **Phaeochroa cuvierii** | 135 |
| **Phaethornis guy** | 157 |
| **Phaethornis longirostris** | 182 |
| **Phaethornis striigularis** | 257 |
| **Phainoptila melanoxantha** | 46 |
| **Phalacrocorax brasilianus** | 239 |
| **Pharomachrus mocinno** | 45 |
| **Pheucticus tibialis** | 40 |
| **Pheugopedius atrogularis** | 72 |
| **Pheugopedius rutilus** | 137 |
| **Philodice bryantae** | 11 |

| | |
|---|---|
| **Mniotilta varia** | 457 |
| **Molothrus aeneus** | 132 |
| **Molothrus bonariensis** | 28 |
| **Molothrus oryzivorus** | 91 |
| **Momotus lessonii** | 312 |
| **Monasa morphoeus** | 30 |
| **Morococcyx erythropygus** | 46 |
| **Morphnarchus princeps** | 58 |
| **Myadestes melanops** | 115 |
| **Mycteria americana** | 252 |
| **Myiarchus crinitus** | 342 |
| **Myiarchus nuttingi** | 71 |
| **Myiarchus panamensis** | 126 |
| **Myiarchus tuberculifer** | 503 |
| **Myiarchus tyrannulus** | 148 |
| **Myiobius atricaudus** | 38 |
| **Myiobius sulphureipygius** | 92 |
| **Myioborus miniatus** | 202 |
| **Myioborus torquatus** | 79 |
| **Myiodynastes hemichrysus** | 43 |
| **Myiodynastes maculatus** | 234 |
| **Myiopagis viridicata** | 93 |
| **Myiophobus fasciatus** | 24 |
| **Myiornis atricapillus** | 36 |
| **Myiothlypis fulvicauda** | 222 |
| **Myiozetetes granadensis** | 289 |
| **Myiozetetes similis** | 844 |
| **Myrmotherula axillaris** | 60 |
| **Myrmotherula schisticolor** | 90 |
| **Notharchus hyperrhynchus** | 108 |
| **Notharchus tectus** | 39 |
| **Nothocercus bonapartei** | 14 |
| **Numenius phaeopus** | 181 |

| | |
|---|---|
| **Philydor rufum** | 10 |
| **Phyllomyias burmeisteri** | 14 |
| **Piaya cayana** | 485 |
| **Piculus simplex** | 56 |
| **Picumnus olivaceus** | 74 |
| **Pionus senilis** | 239 |
| **Piranga bidentata** | 83 |
| **Piranga flava** | 59 |
| **Piranga leucoptera** | 26 |
| **Pitangus sulphuratus** | 878 |
| **Platalea ajaja** | 79 |
| **Platyrinchus coronatus** | 28 |
| **Platyrinchus mystaceus** | 55 |
| **Poecilotriccus sylvia** | 61 |
| **Poliocrania exsul** | 205 |
| **Polioptila plumbea** | 280 |
| **Porphyrio martinica** | 128 |
| **Premnoplex brunnescens** | 61 |
| **Procnias tricarunculatus** | 57 |
| **Progne chalybea** | 371 |
| **Psarocolius montezuma** | 311 |
| **Psarocolius wagleri** | 139 |
| **Pseudastur albicollis** | 86 |
| **Pseudocolaptes lawrencii** | 26 |
| **Psilorhinus morio** | 276 |
| **Psittacara finschi** | 283 |
| **Psittaciformes sp** | 32 |
| **Pteroglossus frantzii** | 95 |
| **Pteroglossus torquatus** | 229 |
| **Ptiliogonys caudatus** | 63 |
| **Pulsatrix perspicillata** | 49 |
| **Pygochelidon cyanoleuca** | 329 |
| **Pyrilia haematotis** | 156 |

| | |
|---|---|
| **Nyctanassa violacea** | 176 |
| **Nyctibius grandis** | 34 |
| **Nyctibius griseus** | 44 |
| **Nycticorax nycticorax** | 70 |
| **Nyctidromus albicollis** | 289 |
| **Odontophorus gujanensis** | 24 |
| **Odontophorus guttatus** | 35 |
| **Odontophorus leucolaemus** | 33 |
| **Oncostoma cinereigulare** | 98 |
| **Onychorhynchus coronatus** | 50 |
| **Oreothlypis gutturalis** | 63 |
| **Ornithion brunneicapillus** | 76 |
| **Ornithion semiflavum** | 35 |
| **Ortalis cinereiceps** | 353 |
| **Oxyruncus cristatus** | 12 |
| **Pachyramphus aglaiae** | 164 |
| **Pachyramphus albogriseus** | 15 |
| **Pachyramphus cinnamomeu** | 173 |
| **Pachyramphus polychopteru** | 115 |
| **Pachyramphus versicolor** | 36 |
| **Pachysylvia decurtata** | 483 |
| **Pandion haliaetus** | 366 |
| **Panterpe insignis** | 42 |
| **Panyptila cayennensis** | 87 |
| **Parabuteo unicinctus** | 23 |
| **Parkesia motacilla** | 118 |
| **Parkesia noveboracensis** | 420 |
| **Passer domesticus** | 138 |
| **Passerina caerulea** | 41 |
| **Passerina ciris** | 63 |
| **Passerina cyanea** | 87 |
| **Patagioenas cayennensis** | 317 |
| **Patagioenas fasciata** | 110 |

| | |
|---|---|
| **Pyrrhura hoffmanni** | 37 |
| **Querula purpurata** | 46 |
| **Quiscalus mexicanus** | 761 |
| **Ramphastos ambiguus** | 318 |
| **Ramphastos sulfuratus** | 385 |
| **Ramphocaenus melanurus** | 187 |
| **Ramphocelus passerinii** | 480 |
| **Ramphocelus sanguinolentu** | 79 |
| **Rhynchocyclus brevirostris** | 63 |
| **Rhytipterna holerythra** | 85 |
| **Rostrhamus sociabilis** | 35 |
| **Rupornis magnirostris** | 381 |
| **Rynchops niger** | 9 |
| **Saltator atriceps** | 106 |
| **Saltator coerulescens** | 191 |
| **Saltator grossus** | 35 |
| **Saltator maximus** | 447 |
| **Saltator striatipectus** | 138 |
| **Sarcoramphus papa** | 119 |
| **Sayornis nigricans** | 128 |
| **Schiffornis veraepacis** | 28 |
| **Sclerurus guatemalensis** | 19 |
| **Sclerurus mexicanus** | 17 |
| **Scytalopus argentifrons** | 69 |
| **Selasphorus flammula** | 37 |
| **Selasphorus scintilla** | 67 |
| **Selenidera spectabilis** | 21 |
| **Semnornis frantzii** | 59 |
| **Serpophaga cinerea** | 38 |
| **Setophaga petechia** | 67 |
| **Setophaga pitiayumi** | 119 |
| **Sipia laemosticta** | 24 |
| **Sittasomus griseicapillus** | 109 |

| | |
|---|---|
| **Patagioenas flavirostris** | 377 |
| **Patagioenas nigrirostris** | 248 |
| **Patagioenas speciosa** | 54 |
| **Patagioenas subvinacea** | 61 |
| **Pelecanus occidentalis** | 351 |
| **Penelope purpurascens** | 251 |
| **Petrochelidon pyrrhonota** | 29 |
| **Peucaea ruficauda** | 134 |
| **Pezopetes capitalis** | 27 |
| **Phaenostictus mcleannani** | 49 |
| **Phaeochroa cuvierii** | 218 |
| **Phaeomyias murina** | 12 |
| **Phaethornis guy** | 223 |
| **Phaethornis longirostris** | 259 |
| **Phaethornis striigularis** | 311 |
| **Phainoptila melanoxantha** | 53 |
| **Phalacrocorax brasilianus** | 346 |
| **Pharomachrus mocinno** | 44 |
| **Pheucticus ludovicianus** | 206 |
| **Pheucticus tibialis** | 55 |
| **Pheugopedius atrogularis** | 75 |
| **Pheugopedius fasciatoventri** | 88 |
| **Pheugopedius rutilus** | 178 |
| **Philodice bryantae** | 46 |
| **Philydor rufum** | 10 |
| **Phyllomyias burmeisteri** | 16 |
| **Phylloscartes superciliaris** | 26 |
| **Piaya cayana** | 664 |
| **Piculus simplex** | 77 |
| **Picumnus olivaceus** | 115 |
| **Pionus menstruus** | 192 |
| **Pionus senilis** | 372 |
| **Piranga bidentata** | 85 |

| | |
|---|---|
| **Spinus psaltria** | 117 |
| **Spinus xanthogastrus** | 33 |
| **Spizaetus ornatus** | 36 |
| **Spizaetus tyrannus** | 56 |
| **Sporophila corvina** | 613 |
| **Sporophila funerea** | 211 |
| **Sporophila minuta** | 64 |
| **Sporophila morelleti** | 279 |
| **Sporophila nigricollis** | 117 |
| **Sporophila nuttingi** | 10 |
| **Sporophila schistacea** | 13 |
| **Stelgidopteryx ruficollis** | 205 |
| **Stelgidopteryx serripennis** | 121 |
| **Stilpnia larvata** | 369 |
| **Streptoprocne rutila** | 38 |
| **Streptoprocne zonaris** | 277 |
| **Sturnella magna** | 141 |
| **Sublegatus arenarum** | 30 |
| **Synallaxis brachyura** | 65 |
| **Syndactyla subalaris** | 20 |
| **Tachybaptus dominicus** | 38 |
| **Tachycineta albilinea** | 164 |
| **Tachyphonus delatrii** | 60 |
| **Tachyphonus luctuosus** | 135 |
| **Tachyphonus rufus** | 77 |
| **Tangara dowii** | 56 |
| **Tangara florida** | 45 |
| **Tangara gyrola** | 145 |
| **Tangara icterocephala** | 179 |
| **Tangara inornata** | 101 |
| **Tangara lavinia** | 17 |
| **Tapera naevia** | 125 |
| **Taraba major** | 61 |

| | |
|---|---|
| **Piranga flava** | 82 |
| **Piranga leucoptera** | 44 |
| **Piranga ludoviciana** | 59 |
| **Piranga olivacea** | 17 |
| **Piranga rubra** | 861 |
| **Pitangus sulphuratus** | 1100 |
| **Platalea ajaja** | 104 |
| **Platyrinchus cancrominus** | 15 |
| **Platyrinchus coronatus** | 52 |
| **Platyrinchus mystaceus** | 60 |
| **Pluvialis squatarola** | 69 |
| **Podilymbus podiceps** | 25 |
| **Poecilotriccus sylvia** | 68 |
| **Poliocrania exsul** | 268 |
| **Polioptila albiloris** | 97 |
| **Polioptila plumbea** | 398 |
| **Porphyrio martinica** | 126 |
| **Porzana carolina** | 12 |
| **Premnoplex brunnescens** | 95 |
| **Procnias tricarunculatus** | 21 |
| **Progne chalybea** | 415 |
| **Protonotaria citrea** | 231 |
| **Psarocolius decumanus** | 84 |
| **Psarocolius montezuma** | 461 |
| **Psarocolius wagleri** | 167 |
| **Pseudastur albicollis** | 145 |
| **Pseudocolaptes lawrencii** | 24 |
| **Psilorhinus morio** | 390 |
| **Psittacara finschi** | 353 |
| **Pteroglossus frantzii** | 118 |
| **Pteroglossus torquatus** | 287 |
| **Ptiliogonys caudatus** | 64 |
| **Pulsatrix perspicillata** | 94 |

| | |
|---|---|
| **Terenotriccus erythrurus** | 38 |
| **Thalasseus maximus** | 50 |
| **Thalurania colombica** | 150 |
| **Thamnistes anabatinus** | 57 |
| **Thamnophilus atrinucha** | 116 |
| **Thamnophilus doliatus** | 269 |
| **Thraupis episcopus** | 871 |
| **Thraupis palmarum** | 490 |
| **Threnetes ruckeri** | 42 |
| **Thripadectes rufobrunneus** | 26 |
| **Thryophilus pleurostictus** | 111 |
| **Thryophilus rufalbus** | 190 |
| **Tiaris olivaceus** | 387 |
| **Tigrisoma fasciatum** | 32 |
| **Tigrisoma lineatum** | 29 |
| **Tigrisoma mexicanum** | 199 |
| **Tinamus major** | 164 |
| **Tityra inquisitor** | 142 |
| **Tityra semifasciata** | 370 |
| **Todirostrum cinereum** | 458 |
| **Todirostrum nigriceps** | 59 |
| **Tolmomyias assimilis** | 43 |
| **Tolmomyias sulphurescens** | 276 |
| **Trochilidae sp** | 85 |
| **Troglodytes aedon** | 722 |
| **Troglodytes ochraceus** | 70 |
| **Troglodytidae sp** | 15 |
| **Trogon caligatus** | 239 |
| **Trogon collaris** | 120 |
| **Trogon massena** | 154 |
| **Trogon melanocephalus** | 148 |
| **Trogon rufus** | 98 |
| **Tunchiornis ochraceiceps** | 66 |

| | |
|---|---|
| **Pygochelidon cyanoleuca** | 377 |
| **Pyrilia haematotis** | 209 |
| **Pyrrhura hoffmanni** | 53 |
| **Querula purpurata** | 61 |
| **Quiscalus mexicanus** | 939 |
| **Ramphastos ambiguus** | 463 |
| **Ramphastos sulfuratus** | 458 |
| **Ramphocaenus melanurus** | 203 |
| **Ramphocelus dimidiatus** | 152 |
| **Ramphocelus passerinii** | 573 |
| **Ramphocelus sanguinolentu** | 87 |
| **Rhodinocichla rosea** | 45 |
| **Rhynchocyclus brevirostris** | 97 |
| **Rhytipterna holerythra** | 110 |
| **Riparia riparia** | 30 |
| **Rostrhamus sociabilis** | 46 |
| **Rupornis magnirostris** | 509 |
| **Saltator atriceps** | 137 |
| **Saltator coerulescens** | 183 |
| **Saltator grossus** | 53 |
| **Saltator maximus** | 530 |
| **Saltator striatipectus** | 126 |
| **Sarcoramphus papa** | 187 |
| **Sayornis nigricans** | 191 |
| **Schiffornis veraepacis** | 54 |
| **Sclerurus guatemalensis** | 20 |
| **Sclerurus mexicanus** | 26 |
| **Scytalopus argentifrons** | 71 |
| **Seiurus aurocapilla** | 74 |
| **Selasphorus flammula** | 46 |
| **Selasphorus scintilla** | 93 |
| **Selenidera spectabilis** | 34 |
| **Semnornis frantzii** | 62 |

| | |
|---|---|
| **Turdus assimilis** | 102 |
| **Turdus grayi** | 946 |
| **Turdus nigrescens** | 38 |
| **Turdus obsoletus** | 32 |
| **Turdus plebejus** | 99 |
| **Tyrannus melancholicus** | 941 |
| **Tyrannus savana** | 103 |
| **Tyto alba** | 24 |
| **Vanellus chilensis** | 219 |
| **Vireo carmioli** | 38 |
| **Vireo flavoviridis** | 350 |
| **Vireo leucophrys** | 51 |
| **Vireolanius pulchellus** | 42 |
| **Volatinia jacarina** | 477 |
| **Xenops minutus** | 146 |
| **Xenops rutilans** | 13 |
| **Xiphorhynchus erythropygiu** | 96 |
| **Xiphorhynchus lachrymosus** | 57 |
| **Xiphorhynchus susurrans** | 269 |
| **Zeledonia coronata** | 24 |
| **Zenaida asiatica** | 345 |
| **Zenaida macroura** | 43 |
| **Zentrygon chiriquensis** | 25 |
| **Zentrygon costaricensis** | 18 |
| **Zentrygon lawrencii** | 13 |
| **Zimmerius parvus** | 260 |
| **Zonotrichia capensis** | 278 |

| | |
|---|---|
| **Serpophaga cinerea** | 55 |
| **Setophaga castanea** | 162 |
| **Setophaga citrina** | 38 |
| **Setophaga coronata** | 44 |
| **Setophaga dominica** | 20 |
| **Setophaga fusca** | 165 |
| **Setophaga magnolia** | 36 |
| **Setophaga pensylvanica** | 693 |
| **Setophaga petechia** | 890 |
| **Setophaga pitiayumi** | 149 |
| **Setophaga ruticilla** | 181 |
| **Setophaga townsendi** | 43 |
| **Setophaga virens** | 167 |
| **Sipia laemosticta** | 47 |
| **Sittasomus griseicapillus** | 136 |
| **Spatula clypeata** | 17 |
| **Spatula discors** | 106 |
| **Sphyrapicus varius** | 40 |
| **Spinus psaltria** | 159 |
| **Spinus xanthogastrus** | 37 |
| **Spiza americana** | 32 |
| **Spizaetus ornatus** | 56 |
| **Spizaetus tyrannus** | 121 |
| **Sporophila corvina** | 637 |
| **Sporophila funerea** | 217 |
| **Sporophila morelleti** | 260 |
| **Sporophila nigricollis** | 58 |
| **Sporophila schistacea** | 16 |
| **Stelgidopteryx ruficollis** | 377 |
| **Stelgidopteryx serripennis** | 276 |
| **Sterna hirundo** | 38 |
| **Stilpnia larvata** | 494 |
| **Streptoprocne rutila** | 51 |

| | |
|---|---|
| **Streptoprocne zonaris** | 314 |
| **Sturnella magna** | 164 |
| **Sublegatus arenarum** | 26 |
| **Sula leucogaster** | 92 |
| **Synallaxis albescens** | 40 |
| **Synallaxis brachyura** | 91 |
| **Syndactyla subalaris** | 39 |
| **Tachybaptus dominicus** | 74 |
| **Tachycineta albilinea** | 306 |
| **Tachyphonus delatrii** | 63 |
| **Tachyphonus luctuosus** | 196 |
| **Tachyphonus rufus** | 100 |
| **Tangara dowii** | 65 |
| **Tangara florida** | 70 |
| **Tangara gyrola** | 241 |
| **Tangara icterocephala** | 277 |
| **Tangara inornata** | 143 |
| **Tangara lavinia** | 33 |
| **Tapera naevia** | 86 |
| **Taraba major** | 84 |
| **Terenotriccus erythrurus** | 92 |
| **Thalasseus elegans** | 43 |
| **Thalasseus maximus** | 231 |
| **Thalasseus sandvicensis** | 129 |
| **Thalurania colombica** | 211 |
| **Thamnistes anabatinus** | 82 |
| **Thamnophilus atrinucha** | 147 |
| **Thamnophilus bridgesi** | 150 |
| **Thamnophilus doliatus** | 326 |
| **Thraupis episcopus** | 963 |
| **Thraupis palmarum** | 626 |
| **Threnetes ruckeri** | 76 |
| **Thripadectes rufobrunneus** | 26 |

| | |
|---|---|
| **Thryophilus pleurostictus** | 101 |
| **Thryophilus rufalbus** | 134 |
| **Tiaris olivaceus** | 439 |
| **Tigrisoma fasciatum** | 54 |
| **Tigrisoma lineatum** | 33 |
| **Tigrisoma mexicanum** | 268 |
| **Tinamus major** | 185 |
| **Tityra inquisitor** | 163 |
| **Tityra semifasciata** | 407 |
| **Todirostrum cinereum** | 528 |
| **Todirostrum nigriceps** | 84 |
| **Tolmomyias assimilis** | 78 |
| **Tolmomyias sulphurescens** | 314 |
| **Touit costaricensis** | 18 |
| **Tringa flavipes** | 62 |
| **Tringa melanoleuca** | 70 |
| **Tringa semipalmata** | 154 |
| **Tringa solitaria** | 110 |
| **Troglodytes aedon** | 731 |
| **Troglodytes ochraceus** | 98 |
| **Trogon bairdii** | 34 |
| **Trogon caligatus** | 290 |
| **Trogon clathratus** | 13 |
| **Trogon collaris** | 151 |
| **Trogon massena** | 236 |
| **Trogon melanocephalus** | 152 |
| **Trogon rufus** | 181 |
| **Tunchiornis ochraceiceps** | 103 |
| **Turdus assimilis** | 92 |
| **Turdus grayi** | 952 |
| **Turdus nigrescens** | 35 |
| **Turdus obsoletus** | 55 |
| **Turdus plebejus** | 146 |

| | |
|---|---|
| **Tyrannulus elatus** | 108 |
| **Tyrannus forficatus** | 165 |
| **Tyrannus melancholicus** | 1229 |
| **Tyrannus savana** | 121 |
| **Tyrannus verticalis** | 50 |
| **Tyto alba** | 21 |
| **Vanellus chilensis** | 209 |
| **Vermivora chrysoptera** | 308 |
| **Vermivora cyanoptera** | 21 |
| **Vireo carmioli** | 54 |
| **Vireo flavifrons** | 519 |
| **Vireo flavoviridis** | 13 |
| **Vireo leucophrys** | 75 |
| **Vireo olivaceus** | 31 |
| **Vireo pallens** | 23 |
| **Vireo philadelphicus** | 379 |
| **Vireo solitarius** | 10 |
| **Vireolanius pulchellus** | 69 |
| **Volatinia jacarina** | 464 |
| **Xenops minutus** | 252 |
| **Xenops rutilans** | 14 |
| **Xiphorhynchus erythropygiu** | 118 |
| **Xiphorhynchus lachrymosus** | 61 |
| **Xiphorhynchus susurrans** | 296 |
| **Zeledonia coronata** | 20 |
| **Zenaida asiatica** | 418 |
| **Zenaida macroura** | 43 |
| **Zentrygon chiriquensis** | 22 |
| **Zentrygon costaricensis** | 12 |
| **Zentrygon lawrencii** | 19 |
| **Zimmerius parvus** | 432 |
| **Zonotrichia capensis** | 326 |

### Appendix 3

Statistics for December Madre Selva dataset

| Threshold | tp | fp | tn | fn | Prediction success | Sorensen index |
|---|---|---|---|---|---|---|
| **100** | 76 | 286 | 289 | 0 | 0.561 | 0.347 |
| **75** | 56 | 23 | 552 | 20 | 0.934 | 0.722 |
| **90** | 65 | 50 | 525 | 11 | 0.906 | 0.680 |

Statistics for June San José dataset

| Threshold | tp | fp | tn | fn | Prediction success | Sorensen index |
|---|---|---|---|---|---|---|
| **100** | 77 | 310 | 119 | 0 | 0.387 | 0.332 |
| **90** | 71 | 158 | 271 | 6 | 0.676 | 0.464 |
| **75** | 66 | 67 | 362 | 11 | 0.846 | 0.628 |
